# Partially observable Markov models inferred using statistical tests reveal context-dependent syllable transitions in Bengalese finch songs

**DOI:** 10.1101/2022.11.15.516592

**Authors:** Jiali Lu, Sumithra Surendralal, Kristofer E. Bouchard, Dezhe Z. Jin

## Abstract

Generative models have broad applications, ranging from language processing to analyzing bird-song. In this study, we demonstrate how a statistical test, designed to prevent overgeneralization in sequence generation, can be used to deduce minimal models for the syllable sequences in Bengalese finch songs. We focus on the partially observable Markov model (POMM), which consists of states and the probabilistic transitions between them. Each state is associated with a specific syllable, with the possibility of multiple states being associated to a single syllable. This feature sets the POMM apart from a standard Markov model, where each syllable is associated to just one state. This multiplicity suggests that syllable transitions are influenced by the specific contexts in which the transitions appear. We apply this method to analyze the songs of six adult male Bengalese finches, both before and after they are deafened. Our findings indicate that auditory feedback is crucial in shaping the context-dependent syllable transitions characteristic of Bengalese finch songs.

**Significance:** Generative models are adept at representing sequences where the order of elements, such as words or birdsong syllables, depends on the context. In this study, we demonstrate that a probabilistic model, inspired by neural encoding of song production in songbirds, effectively captures context-dependent transitions of syllables in Bengalese finch songs. Our findings indicate that the absence of auditory input, as seen in deafened finches, diminishes these context dependencies. This implies that auditory feedback is vital for establishing context-based sequencing in their songs. Our method can be applied to various behavioral sequences, offering insights into the neural underpinnings that govern statistical patterns in these sequences.

## Introduction

Behavioral sequences, ranging from human language to birdsong, follow probabilistic rules. The success of large language models, such as GPT, demonstrates that the transition probabilities between words in language are highly influenced by the preceding words (OpenAI, 2023). However, it remains unclear how such context-dependent probabilistic rules are encoded in the brain. Studies of birdsongs provide insights into this issue. Variable songs of species such as the Bengalese finch and canary exhibit context-dependent syllable transitions (Okanoya, 2004; Jin and Kozhevnikov, 2011; Jin, 2013; Markowitz et al., 2013), and the neural correlates of these dependencies have been studied using advanced imaging techniques to visualize the neural activities in the brain areas controlling the song (Cohen et al., 2020).

The probabilistic rules of syllable transitions, or the song syntax, can be effectively modeled using state transition models. This approach has been detailed in several studies (Okanoya, 2004; Jin and Kozhevnikov, 2011; Jin, 2013; Markowitz et al., 2013). Specifically, the syllable sequences in Bengalese finch songs, excluding syllable repetitions, are effectively characterized by Partially Observable Markov Models (POMMs) (Jin and Kozhevnikov, 2011).

A POMM consists of a number of states, each associated with a syllable, along with a start state and an end state. There are fixed probabilities dictating the transitions from one state to another. Beginning from the start state, a sequence of states is generated via these transitions until the end state is reached. This state sequence then determines the corresponding syllable sequence through the established state-syllable associations. Importantly, the transition probabilities from any given state are independent of the path taken to reach that state. Consequently, the dynamics of state transitions within a POMM are Markovian. In a POMM, a single syllable may be associated with multiple states. This feature of state multiplicity enables the Markovian dynamics of state transitions to produce non-Markovian syllable sequences. As a result, syllable transitions can become context-dependent (Jin and Kozhevnikov, 2011).

The POMM framework draws inspiration from the hypothesis that the generation of birdsong is driven by synaptic chain networks within HVC, a key premotor nucleus in the song system (Nottebohm et al., 1976). This concept has been supported by a number of studies (Hahnloser et al., 2002; Fee et al., 2004; Jun and Jin, 2007; Jin et al., 2007; Jin, 2009; Long et al., 2010; Wittenbach et al., 2015; Lynch et al., 2016; Picardo et al., 2016; Jin, 2013; Zhang et al., 2017; Egger et al., 2020; Tupikov and Jin, 2021). More specifically, the hypothesis posits that neurons within the HVC, which projects to downstream motor areas, are organized into feedforward synaptic chains (Jin et al., 2007; Long et al., 2010; Egger et al., 2020; Tupikov and Jin, 2021). During the production of a syllable, there is a sequential activity propagating along a chain, where each neuron discharges a single burst (Hahnloser et al., 2002; Lynch et al., 2016; Fee et al., 2004; Jin, 2009). Such a “syllable-chain” can be conceptualized as a specific state within a POMM (Jin, 2009; Jin and Kozhevnikov, 2011; Wittenbach et al., 2015). Utilizing POMM to analyze observed sequences of birdsong syllables offers a unique perspective, potentially unveiling the intricate neural dynamics within the HVC that facilitate the creation of variable syllable sequences (Jin, 2009; Wittenbach et al., 2015).

Auditory feedback plays a crucial role in shaping the syllable sequences of Bengalese finch songs (Okanoya and Yamaguchi, 1997; Woolley and Rubel, 1997; Woolley and Rubel, 2002; Sakata and Brainard, 2008; Wittenbach et al., 2015). Within a few days after deafening, the syllable sequences become more random (Okanoya and Yamaguchi, 1997; Woolley and Rubel, 1997). Additionally, there is a significant decrease in syllable repetitions (Wittenbach et al., 2015). When altered auditory feedback is provided to Bengalese finches during singing, particularly at the branching points of syllable transitions, it can significantly influence the probabilities of these transitions (Sakata and Brainard, 2006; Sakata and Brainard, 2008). These observations indicate that auditory feedback may play a significant role in shaping the context dependencies of syllable transitions in the song of the Bengalese finch.

In this paper, we analyze context dependencies in the songs of six Bengalese finches before and shortly after deafening. We first develop a novel method for inferring a POMM from a set of observed syllable sequences. This method is based on the concept of “sequence completeness,” denoted as *P*_*c*_, which is the cumulative probability of the POMM generating each unique sequence in the observed set. Additionally, we incorporate the differences between the probabilities of unique sequences as calculated from the observed set and those predicted by the model, leading to an “augmented sequence completeness,” denoted as *P*_*β*_. Our method aims to identify the minimum number of states for each syllable necessary to achieve statistical compatibility between the POMM and the observed sequences. This is accomplished by testing the hypothesis that the observed sequences were generated by the POMM, using the distribution of *P*_*β*_ for sets of sequences generated by the POMM.

Our new method for inferring a POMM from syllable sequences in Bengalese finch songs presents a significant advancement over the previous heuristic approach (Jin and Kozhevnikov, 2011). The earlier method involved initially constructing a tree POMM capable of precisely replicating the observed sequences. This was then followed by a process of simplification, which primarily consisted of merging states to minimize the number of states. The effectiveness of the mergers was evaluated using various statistical measures, such as n-gram distributions (which assess the probabilities of observing subsequences of length *n*) and the probabilities of encountering specific syllables at different points in the sequence. This process required manual intervention, adding a layer of complexity (Jin and Kozhevnikov, 2011). In contrast, our new method is fully automated, eliminating the need for manual adjustments. Moreover, it relies on a single, interpretable statistical measure, simplifying the process of model induction. The simplicity and principled nature of the approach potentially make our new method more accessible and robust in studying behavioral sequences.

Using this novel method, we demonstrate that auditory feedback plays a crucial role in shaping the context dependencies in the songs of Bengalese finches. We achieve this by inferring minimal POMMs for the syllable sequences of six Bengalese finches, both under normal conditions and shortly after deafening. In the normal condition, we find that the POMMs for all the birds necessitate multiple states for certain syllables, indicating complex context dependencies. However, shortly after the birds are deafened, there is a significant shift in the structure of these POMMs. For half of the birds (3 out of 6), the POMMs essentially simplify to standard Markov models, indicating a loss of context-dependent syllable transitions. For the remaining birds (also 3 out of 6), there is a substantial reduction in state multiplicity within the POMMs. These observations show that the loss of auditory feedback weakens the context dependencies in the syllable transitions of Bengalese finch songs, underscoring the importance of hearing in maintaining the complexity of their song structure.

## Results

### The dataset

In our previous study (Wittenbach et al., 2015), we focused on the phenomenon of syllable repetition in the songs of six Bengalese finches, particularly examining the role of auditory feedback. Our findings suggest that long sequences of repeated syllables are best described by a non-Markovian process (Jin and Kozhevnikov, 2011). The process is characterized by a gradual decrease in the probability of a syllable repeating itself as the sequence progresses. We identified the underlying neural mechanism as the synaptic adaptation in the auditory feedback pathway. Essentially, strong and adapting auditory feedback is crucial for sustaining long syllable repetitions. Supporting this, we observed that removing auditory feedback through deafening led to a significant shortening of these syllable repetitions (Wittenbach et al., 2015). It is important to note, however, that our analysis was specifically confined to syllable repetitions and did not extend to other aspects of the song structure.

In this work, we study the context dependencies in the syllable sequences of the six Bengalese finches and how auditory feedback might contribute to them using the POMMs. Because syllable repetitions are best described by non-Markovian state transition models, we focus on the non-repeat versions of the sequences, in which only the first syllable of any repetition is retained. For example, if the syllable sequence is *ABBBC*, the non-repeat version is *ABC*. In the rest of the paper, the term “syllable sequences” specifically refers to these non-repeat versions. Each syllable sequence is typically preceded by a variable number of introductory notes, which are excluded from the analysis.

### POMM diagram

POMMs are visualized with directed graphs (Fig. 1). Following the convention introduced previously (Jin and Kozhevnikov, 2011), we denote the start state as a pink node marked with the symbol *α*. All other states are represented as nodes marked with associated syllables. Syllables with multiple states are marked in red fonts. A state’s color is cyan if it can transition to the end state, and white otherwise. To reduce clutter in the graph, the end state is not shown. When necessary, we use the symbol *ω* to mark the end of sequences. State transitions are shown with arrows color-coded for the transition probabilities *P*. Strong transitions (0.5 *≤ P ≤* 1) are shown in red; medium transitions (0.1 *≤ P <* 0.5) in green; and weak transitions (0.01 *≤ P <* 0.1) in gray. To further reduce clutter, only transitions with *P ≥* 0.01 are shown.

**Figure 1.**
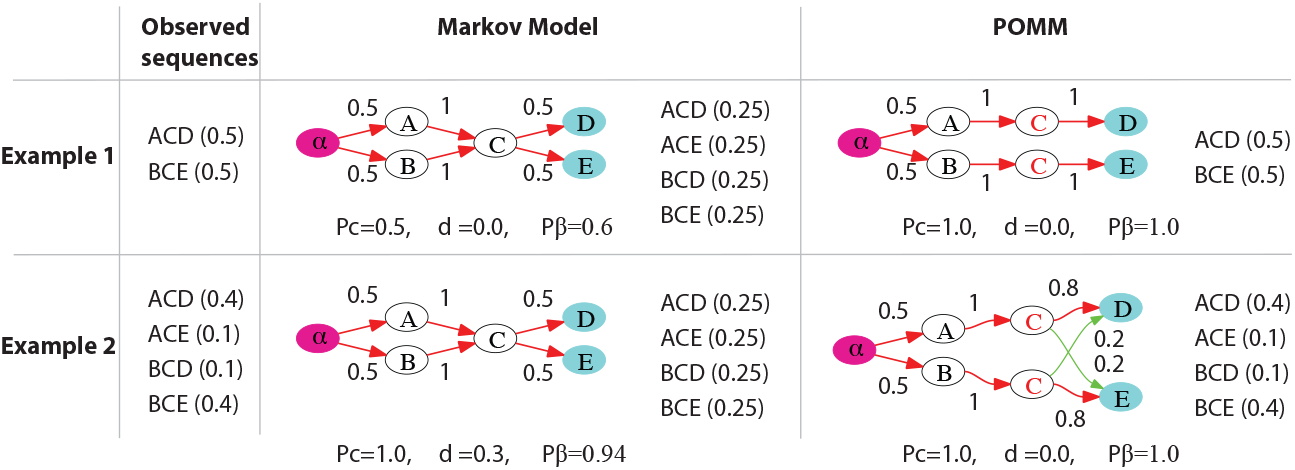
Two examples illustrating two types of context dependency. In Example 1, there are two unique sequences with equal probabilities (shown in parentheses) in the observed set. The Markov model overgeneralizes, resulting in two unobserved sequences and a sequence completeness of *P*_*c*_ = 0.5. The POMM with two states for syllable *C* avoids overgeneralization, achieving *P*_*c*_ = 1 and *P*_*β*_ = 1. In Example 2, the observed set comprises four unique sequences, with two being more frequent. The Markov model fails to capture these frequency differences (total variation distance *d* = 0.3), even though *P*_*c*_ = 1. However, the POMM with two states for syllable *C* successfully captures the probabilities of the unique sequences, indicated by *d* = 0 and *P*_*β*_ = 1.

### Two types of context dependency

We construct two simple examples to demonstrate the existence of different types of context dependencies in syllable transitions (Fig. 1). The examples contain five syllables, *A, B, C, D, E*. The transitions from *C* to *D* or to *E* depend on whether *C* is preceded by *A* or *B*.

Example 1 demonstrates a case where a syllable transition can be allowed or prohibited depending on the context. The observed sequences contain two unique sequences: *ACD* and *BCE*, each with a probability of 0.5. The transition *C → D* occurs only if *C* is preceded by *A*, and the transition *C → E* occurs only if *C* is preceded by *B*. Consequently, sequences *ACE* and *BCD* are not observed. We refer to this form of context dependence as *type I context dependence*.

Example 2 demonstrates a case where context dependence manifests in the transition probabilities. The observed sequences contain four unique sequences: *ACD*, with a probability of 0.4; *ACE*, with a probability of 0.1; *BCD*, with a probability of 0.1; and *BCE*, with a probability of 0.4. The transitions *C → D* and *C → E* are observed regardless of the preceding syllable. However, the transition probabilities differ depending on whether *A* or *B* precedes *C*. We refer to this kind of context dependence as *type II context dependence*.

A simple model that can be inferred from the observed sequences is the Markov model, in which all syllables are associated with one state. For both examples, the Markov model is too simple to capture the context dependencies.

The Markov model for Example 1 can be inferred by calculating transition probabilities from the observed sequences (Fig. 1). The sequences can start with either *A* or *B* with equal probability, hence the start state transitions to the states associated with *A* or *B* (*A*-state or *B*-state) with probability 0.5. These two states transition to the *C*-state with probability 1. Since *C* can be followed by either *D* or *E*, the *C*-state transitions to the *D*-state or the *E*-state, each with a probability of 0.5. Finally, the *D*-state and the *E*-state transition to the end state with probability 1.

The Markov model fails because it overgeneralizes. From the start state, there are four possible state transition paths, generating four sequences *ACD, ACE, BCD*, and *BCE*, each with a probability of 0.25. However, the sequences *ACE* and *BCD* are not observed in the data.

The issue of overgeneralization can be quantified using the concept of *sequence completeness P*_*c*_, which is defined as the total probability of the model generating all unique sequences in the observed set of sequences:

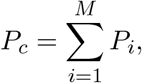

where *M* is the number of unique sequences, and *P*_*i*_ is the probability of the *i*-th unique sequence computed with the model. The degree of overgeneralization is represented by 1 *− P*_*c*_, which corresponds to the total probability of generating sequences with the model that are not present in the observed set. For Example 1, the unique sequences in the data are *ACD* and *BCE*. The probabilities of the Markov model generating these sequences are 0.25 each. Therefore, we calculate *P*_*c*_ = 0.5.

To avoid overgeneralization, we need to infer a more complex model, a POMM in which *C* is associated with two states, with the *A*-state and the *B*-state transitioning separately to each of these states (Fig. 1). This POMM generates the two sequences *ACD* and *BCE*, each with a probability of 0.5, resulting in *P*_*c*_ = 1 for the observed set. In other words, the model does not overgeneralize.

In Example 2, the Markov model fails not due to overgeneralization but because the probabilities do not align with the data (Fig. 1). The model generates all observed unique sequences, hence *P*_*c*_ = 1. However, the less frequent sequences *ACE* and *BCD*, with a probability of 0.1 in the data, are assigned a higher probability in the Markov model (0.25 for each). Conversely, the more probable sequences *ACD* and *BCE*, with a probability of 0.4 in the data, are represented as less frequent in the model, each with a probability of 0.25.

A simple measure of the differences in probabilities is the total variation distance (Gibbs and Su, 2002), defined as

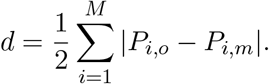

Here,

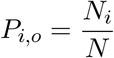

represents the observed probability of the *i*-th unique sequence. It is the ratio of the copy number *N*_*i*_ of this sequence to the total number *N* of sequences observed; and *P*_*i,m*_ is the normalized probability of the sequence computed with the model, given by

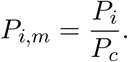

The normalization is to ensure that

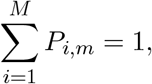

which is necessary since we are comparing *P*_*i,m*_ with *P*_*i,o*_, where 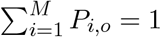.

In Example 2, the Markov model has *d* = 0.3. A more complex model, which includes two states for *C* as shown in Fig. 1, has *d* = 0. This indicates that it effectively captures the context-dependent changes in transition probabilities.

The total variation distance may not adequately reveal type I context dependence. In Example 1, the Markov model generates the two observed sequences *ACD* and *BCE* with probabilities of 0.25 each. However, after normalization, these probabilities become 0.5. Consequently, for the Markov model, we have *d* = 0. Thus, a perfect match in probabilities as indicated by the total variation distance does not necessarily guarantee that the model is accurate.

To capture both type I and type II context dependencies, we combine *P*_*c*_ and *d* into a single measure

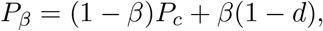

where *β* is the weight given to the total variation distance, and is a number between 0 and 1. A perfect model would have *P*_*β*_ = 1. We refer to this combined measure as the *augmented sequence completeness*.

As we will demonstrate, model selection based on *P*_*β*_ is not sensitive to the value of *β* as long as it is neither close to 0 nor 1. A suitable choice of *β* should balance the variances of *P*_*c*_ and *d* to ensure that the variance of *P*_*β*_ is not predominantly influenced by either. Since *P*_*c*_ is a sum of probabilities, it tends to be less sensitive to measurement errors, as positive and negative errors often offset each other. Conversely, *d* aggregates the absolute values of errors and is likely more sensitive to measurement inaccuracies. Therefore, we should choose *β <* 0.5 to reflect this difference in variations. In this study, we set *β* = 0.2. This consideration becomes crucial when the number of observed sequences is small. With a large number of sequences, accurate probability measurements are expected, rendering the specific choice of *β* less critical.

The POMMs with two states for *C* in both Example 1 and Example 2 yield *P*_*β*_ = 1, irrespective of the chosen value for *β*.

### Inference of a minimal POMM from observed sequences

In this section, we demonstrate how POMMs can be inferred from syllable sequences using a statistical test. For this purpose, we construct a “ground-truth” POMM, from which we generate “observed sequences” (Fig. 2). We then apply the inference process to these observed sequences to infer minimal POMMs. Finally, we compare the inferred POMMs with the original ground-truth POMMs for validation.

**Figure 2.**
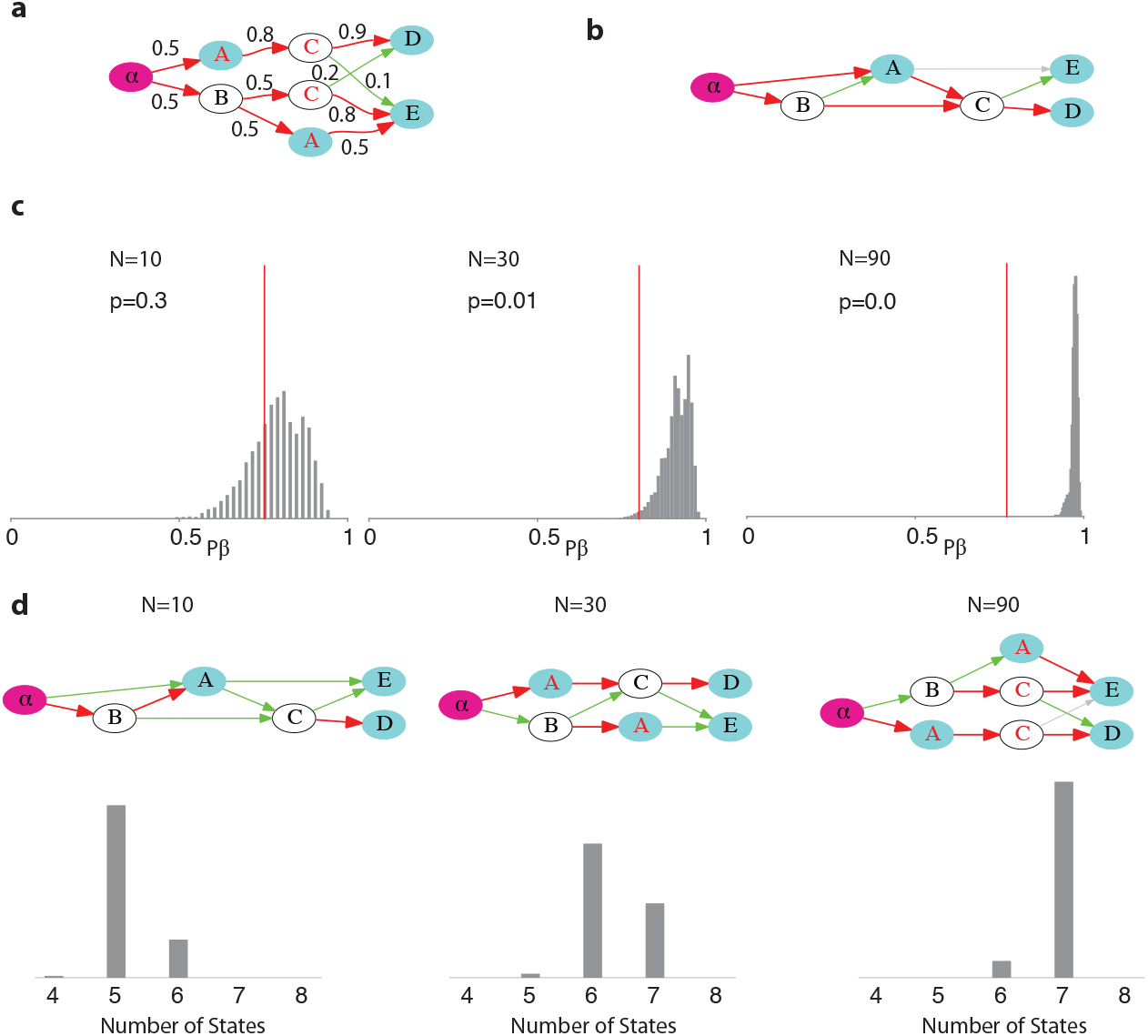
An example for statistically testing and inferring POMMs from a finite set of observed sequences. **a**. The ground truth POMM from which the observed sequences are generated. The numbers near the arrows are the transition probabilities. **b**. The Markov model for a set of *N* = 30 observed sequences. **c**. Statistical tests of the Markov models for *N* = 10, 30, 90. The gray bars are the distributions of *P*_*β*_ for the generated sets. The red lines are the *P*_*β*_ for the observed sets. **d**. POMMs inferred from the observed sets, and distributions of the number of states in the POMMs for 100 runs.

#### Statistical test of a POMM

To identify a POMM that aligns with the observed set of syllable sequences, it is necessary to develop a method for statistically evaluating the POMM’s fit. The evaluation can be cast as a hypothesis test, where the null hypothesis posits that the observed set is generated by the POMM. *P*_*β*_ can be utilized for this evaluation. Ideally, *P*_*β*_ of the observed set, as computed with the POMM, should be 1. This indicates that the POMM accurately generates all the unique sequences present in the observed set and, crucially, does not produce unobserved sequences. Moreover, it suggests that the probabilities of these unique sequences are consistent with the observations. However, in practice, due to the finite number *N* of observed sequences, the observed set might not encompass all sequences a bird is capable of producing. Therefore, a *P*_*c*_ value less than 1 could stem from the limited size of *N*, rather than from the model’s overgeneralization. Furthermore, discrepancies in the probabilities of the unique sequences might arise from imprecise probability measurements when *N* is finite.

To account for the finite *N* effect, we generate random sets of *N* sequences from the POMM. For each generated set, we compute *P*_*β*_ using the POMM. The distribution of *P*_*β*_ for these generated sets helps assess the likelihood that the *P*_*β*_ of the observed set is part of this distribution. Specifically, we calculate the probability *p* that the observed *P*_*β*_ exceeds the *P*_*β*_ of the generated sets. If *p <* 0.05, we infer that the observed *P*_*β*_ is unlikely to have been drawn from this distribution, leading to the rejection of the POMM as the model for the observed set. Conversely, if *p ≥* 0.05, the POMM is not statistically rejected and is therefore accepted as the model for the observed set. To construct the *P*_*β*_ distribution, we generate 10,000 random sets of *N* sequences from the POMM.

To demonstrate this process, we use the “ground truth model” shown in Figure 2a. This model comprises two states for syllables *A* and *C*, and one state for each of the syllables *B, D*, and *E*. The model is capable of generating seven unique sequences with associated probabilities: *A* with a probability of 0.1; *ACD* with a probability of 0.36; *ACE* with a probability of 0.04; *BCD* with a probability of 0.05; *BCE* with a probability of 0.2; *BAE* with a probability of 0.125; and *BA* with a probability of 0.125. The sequences produced by the ground truth model include both type I and type II context-dependent syllable transitions.

The ground-truth model is evidently non-Markovian. Ideally, when *N* is large, the Markov model inferred from the observed sequences should be rejected by the statistical test with *P*_*β*_. To demonstrate this, we generate three sets of observed sequences from the ground truth model, using *N* = 10, *N* = 30, and *N* = 90. For each set, we infer the Markov models and subsequently test their fit.

Markov models are inferred by analyzing observed sequences. We count the number of transitions between syllables, as well as transitions from the start to the syllables and from the syllables to the end. These counts are then converted into transition probabilities through normalization. The Markov model inferred from the set with *N* = 30 is illustrated in Fig. 2b. To assess the validity of a Markov model, we generate 10,000 random sets of *N* sequences using the model. For each set, we compute *P*_*β*_. This computation yields a distribution of *P*_*β*_, as shown in Fig. 2c. We then compare this distribution to the *P*_*β*_ of the observed set.

As *N* increases, the distribution of *P*_*β*_ shifts towards 1. This shift occurs because a larger *N* ensures that most unique sequences the model can generate are included in a generated set, leading to *P*_*c*_ *→* 1 for these sets. Furthermore, the probabilities of the unique sequences computed with the generated sets align more closely with those computed using the model. However, the *P*_*β*_ of the observed sets, as indicated by red lines in the figure, remains relatively unchanged with increasing *N*. This stability is attributed to the fact that the observed sets are generated using the ground truth model, which is not a Markov model. Therefore, increasing *N* does not enhance *P*_*c*_ for the observed sets. As *N* grows, the *P*_*β*_ for the observed sets consistently falls below that of the generated sets, suggesting that the Markov models are not statistically compatible with the observed sets.

In the examples depicted in the figure, the p-values are *p* = 0.3 for *N* = 10, *p* = 0.01 for *N* = 30, and *p* = 0 for *N* = 90. The results may exhibit variability due to differences in the samples of the “observed sequences” generated from the ground truth model. To demonstrate these fluctuations, we repeated the previously described process 100 times for each *N*. For *N* = 10, the results were *p* = 0.3 *±* 0.3 (mean *±* standard deviation); for *N* = 30, *p* = 0.01 *±* 0.03; and for *N* = 90, *p* = 0 *±* 0. Consequently, using the *p <* 0.05 criterion, the Markov model is rejectable for *N* = 30 and *N* = 90. However, for *N* = 10, the model cannot be rejected, even though the ground truth model is non-Markovian. This occurs because, when *N* is too small, there is insufficient evidence to reject the Markov model.

If the ground truth model is Markovian, an increase in *N* does not result in the rejection of the Markov model, which aligns with expectations (Supplementary Fig. S1).

While the Markov model serves as an example, the process of statistical testing based on *P*_*β*_ is applicable for evaluating any POMM.

#### Searching the state space for a minimal POMM

The statistical test described above enables the identification of POMMs compatible with a specific set of observed sequences. Additionally, we require that the POMM must represent a minimal model, characterized by the smallest possible number of states for each syllable. This objective is accomplished by exploring the state space of the POMMs. Our initial step involves identifying a POMM that meets the criteria of the statistical test. Subsequently, we simplify this model by merging and deleting states until the POMM is reduced to the simplest form that still satisfies the statistical test requirements.

To search for a POMM compatible with the observed set, we construct higher-order Markov models (Supplementary Fig. S2). To derive the *m*-th order Markov model, we begin by flanking each sequence in the observed set with the start symbol *α* and the end symbol *ω*. We then collect unique subsequences of length *m*, along with subsequences up to length *m* starting from *α*. For instance, with *m* = 2, the sequence *αACDω* yields the unique subsequences *α, αA, AC, CD*, and *Dω*. Each unique subsequence is assigned to a state. The subsequence *α* is assigned to the start state, while all subsequences ending with *ω* are assigned to the end state. The remaining unique subsequences are each assigned to a distinct state, with the final syllable as the symbol for the state. This assignment transforms each observed sequence into a sequence of states. We then calculate the transition probabilities between these states by tallying the number of transitions. This method produces a POMM equivalent to the *m*-th order Markov model. We apply the statistical test to ascertain if the POMM meets the acceptance criterion of *p ≥* 0.05. Starting from *m* = 1, which represents the basic Markov model, we incrementally increase *m* until the POMM is accepted.

We next reduce the POMM by merging and deleting states associated with the same syllable. The state transition probabilities are re-computed using the counts of transitions between the states (Jin and Kozhevnikov, 2011). A merge is kept if the resulting POMM is accepted. After the merging stops, we further reduce the POMM through state deletion. If a syllable is associated with more than one state, we reduce the number of states for that syllable by 1. We compute the transition probabilities between the states by maximizing the log-likelihood that the model generates the observed sequences using the Baum-Welch algorithm (Rabiner, 1989). To avoid local minima that the algorithm may encounter, we consider 100 runs of the algorithm with random seeds and select the run with the highest likelihood. If the reduced POMM is accepted, deletion continues. With these state reduction procedures, we arrive at a POMM with the minimal number of states that passes the statistical test based on *P*_*β*_.

After state reduction, we simplify the transitions between the states in the POMM. We systematically cut every transition and recalculate the transition probabilities using the Baum-Welch algorithm. If the log-likelihood of the observed sequences after the cut is larger than a threshold, the cut is accepted; otherwise the transition is retained. The threshold is set to the log-likelihood before the cuts minus an estimate of the fluctuation of the log-likelihood due to inaccuracies in computing the transition probabilities. The estimate is set to be the standard deviation of the log-likelihoods before the cuts in the 100 runs of the Baum-Welch algorithm with random seeds. If the POMM after the cuts does not pass the statistical test, the cuts are reversed.

#### Evaluation with the ground truth model

We evaluate the aforementioned procedure using the ground truth model (Fig. 2a). A set of *N* “observed sequences” is generated from the ground truth model, from which we infer the minimal POMMs. To assess the impact of sampling, this process is repeated 100 times. The results for *N* = 10, 30, 90 are depicted in Fig. 2d. We present typical POMMs inferred, along with the distributions of the total number of states in the POMMs inferred from the 100 sets. For *N* = 10, the total number of states is predominantly 5, and the Markov model is generally accepted. Some models have 4 states. This is because syllables *D* or *E* may not appear in the observed sequences due to the small sample size *N*. When *N* = 30, the total number of states varies between 5 to 7, with a typical POMM having 6 states, as illustrated in the figure. For *N* = 90, the total number of states is primarily 7, and the inferred POMMs closely resemble the structure of the ground truth model. As we have discussed earlier, the results are insensitive to the choice of *β* (Supplementary Fig. S3).

This example demonstrates that our procedure tends to infer a simpler POMM than the ground truth model when *N* is small. Conversely, when *N* is large, the procedure successfully uncovers the ground truth model. Notably, the procedure does not generate models that are more complex than the ground truth model.

### POMMs of Bengalese finch songs in the normal condition and after deafening

To investigate the effect of auditory input on the song syntax of the Bengalese finch, we analyzed the songs of six adult Bengalese finches before and two days after deafening. This dataset has been previously utilized for analyzing syllable repeats (Wittenbach et al., 2015). In this study, we focus on the non-repetitive versions of the syllable sequences.

#### Test of Markov models on the Bengalese finch songs

We tested whether Markov models are statistically compatible with the observed syllable sequences, using the *p ≥* 0.05 criterion. The results are presented in Supplementary Figs. S4-S5. For three birds, the syllable sequences are incompatible with the Markov models both before and after deafening (o10bk90, normal *p* = 0, deafened *p* = 0; bfa16, normal *p* = 0, deafened *p* = 0; o46bk78, normal *p* = 0, deafened *p* = 0). For the other three birds, the sequences are incompatible with the Markov models before deafening, but post-deafening, the Markov models are accepted (bfa7, normal *p* = 0, deafened *p* = 0.4; bfa14, normal *p* = 0, deafened *p* = 0.5; bfa19, normal *p* = 0.03, deafened *p* = 0.3). These results suggest that deafening may reduce the Bengalese finch song syntax from non-Markovian to Markovian for some birds, but not universally.

Deafening results in the creation of novel transitions between syllables, as well as new starting and ending syllables. The transition probabilities for these novel transitions tend to be low (median, 0.04), yet 22% exceed a probability of 0.1 (representing 18 out of 81 transitions). The majority of these novel transitions are observed in two birds (27 for bfa14; 21 for bfa19). Additionally, a small number (8) of transitions disappear following deafening, with a median transition probability in the normal condition of 0.02.

As observed in previous studies (Woolley and Rubel, 1997; Okanoya and Yamaguchi, 1997), deafening increases sequence variability. The variability of transitions from a given syllable *i* (or the start state) is quantified with the transition entropy 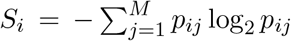, where *M* represents the number of branches of the transitions, and *p*_*ij*_ is the probability of the *j*-th branch. If *M* = 1, the transition is stereotypical and *S*_*i*_ equals zero. For a given *M*, the entropy reaches its maximum when the transition probabilities for all branches are equal. This maximum entropy increases with *M*. The median of transition entropies is significantly higher after deafening (0.95 *±* 0.55) than before (0.35 *±* 0.51; *p* = 5 *×* 10^*−*6^, Wilcoxon signed-rank one-sided test). Similarly, the number of branches *M* is also significantly larger after deafening (4 *±* 1.5, median *±* s.d.) than before (2 *±* 0.90; *p* = 9.8 *×* 10^*−*7^, Wilcoxon signed-rank one-sided test).

#### POMMs of the Bengalese finch songs

We inferred minimal POMMs from the observed syllable sequences before and after deafening in six birds (Figs. 3-4). In the normal condition, the songs of the birds comprise 44 syllables; of these, 26 require 1 state, 13 require 2 states, 2 require 3 states, and 3 require 4 states. Thus, most syllables necessitate 1 or 2 states. The POMMs encompass 76 states in total. When considering only transition branches with probabilities greater than 0.01, the majority of states have up to 3 outgoing branches, with 29, 31, and 11 states having 1, 2, and 3 branches, respectively. After deafening, there are 43 syllables (syllable *g* for bfa7 drops out after deafening). Most syllables (40) require only 1 state, and the remaining 3 require 2 states. There are 52 states in the POMMs. Counting only the transition branches with probabilities greater than 0.01, the states have up to 7 outgoing branches (there are 2, 19, 7, 13, 6, 3, 2 states with branch numbers 1 to 7, respectively).

**Figure 3.**
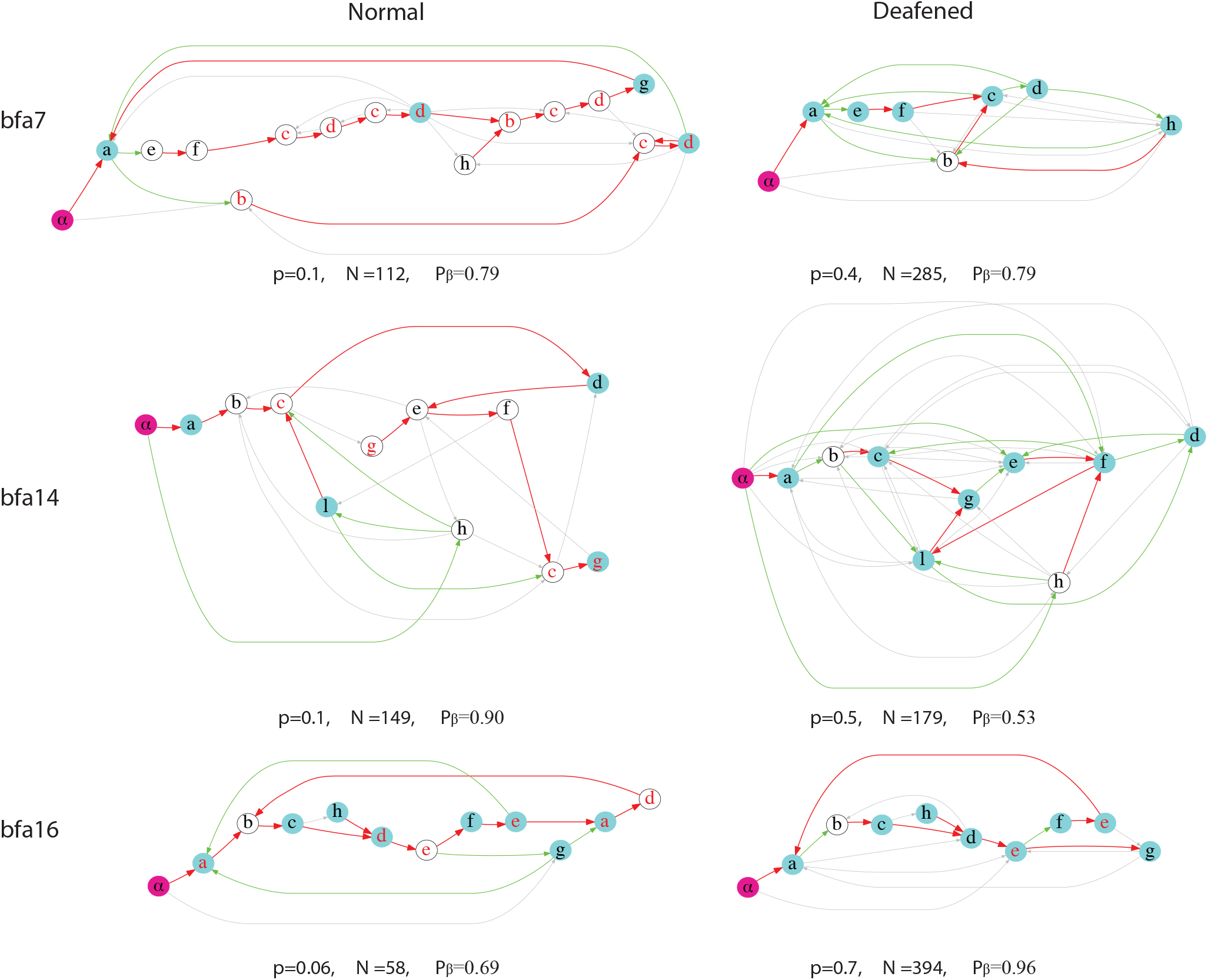
POMMs before and after deafening for three birds. The results for bfa7, bfa14, and bfa16 are shown. The p-values, the number of sequences in the observed sets, and *P*_*β*_ of the observed sets are shown.

**Figure 4.**
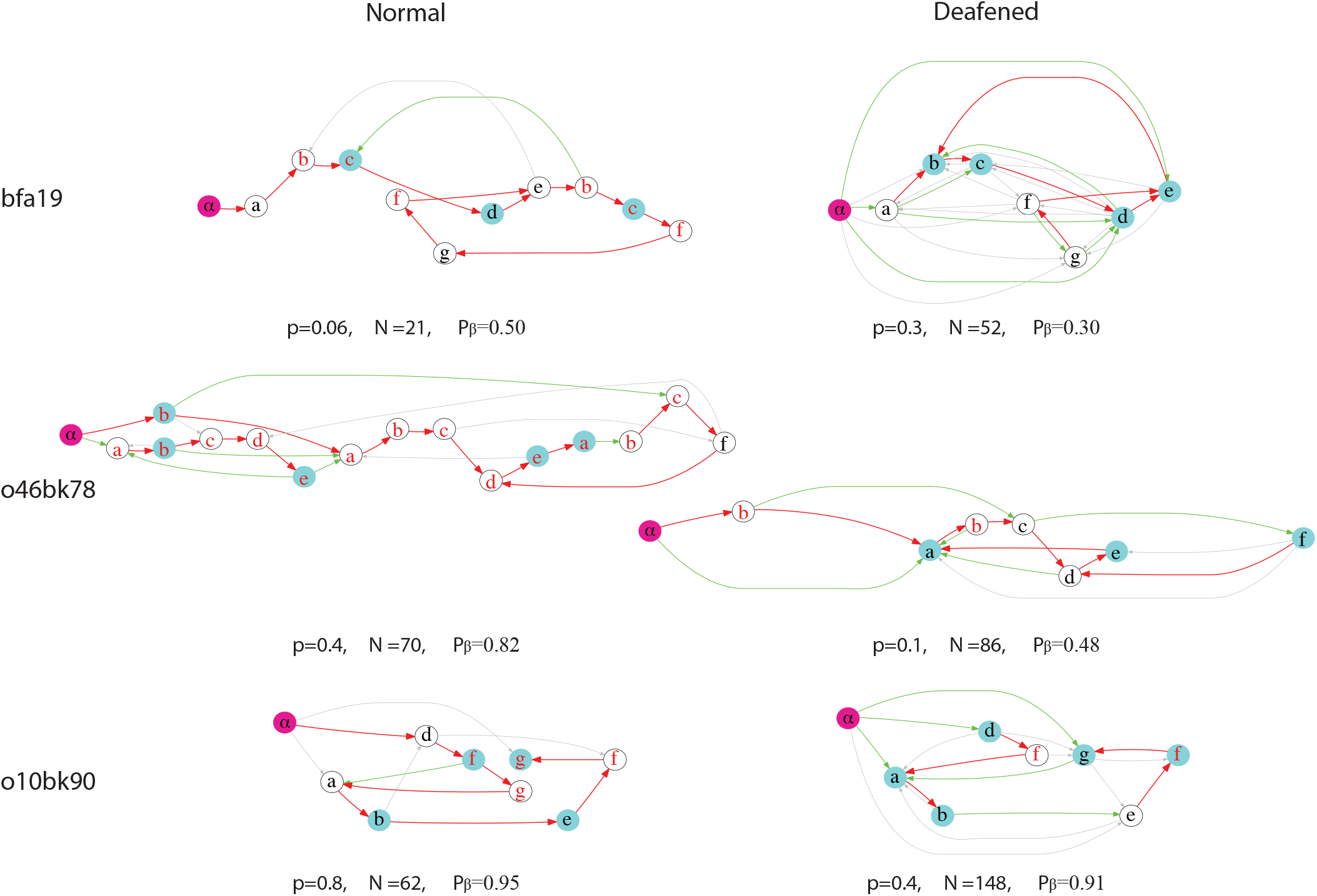
POMMs before and after deafening for the other three birds. The results for bfa19, ok46bk78, and o10bk90 are shown.

**Figure 5.**
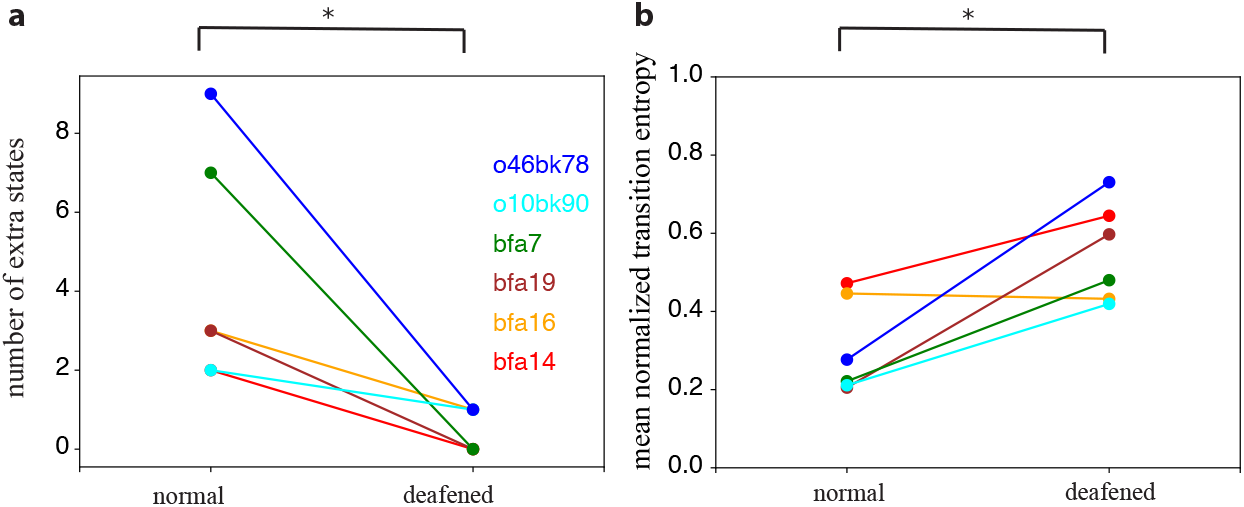
Effects of deafening on the POMMs. **a**. The numbers of extra states for the syllables are significantly reduced. **b**. The transitions from the states become more random after deafening.

Deafening significantly reduces state multiplicity, as evidenced by the decrease in the number of extra states (defined as the number of states for the syllables minus the number of syllables) (Fig.5a; Wilcoxon signed-rank one-sided test, *p* = 0.02). The mean normalized transition entropy between states, which is the transition entropy divided by log_2_ *M*, is notably higher after deafening in all but one bird (Fig.5b; Wilcoxon signed-rank one-sided test, *p* = 0.03, tested across all birds). Therefore, deafening reduces context dependencies in syllable sequences, as demonstrated by the diminished state multiplicity. Furthermore, the transitions between states become more random.

The POMMs encode context dependencies in the syllable transitions. Detailed analysis for the individual birds can be found in the Supporting Information.

### Neural mechanisms of the state multiplicity

Within the framework of syllable-chains, syllables are driven by chain networks in the HVC. The propagation of spikes from one chain to another results in syllable transitions. The intrinsic model provides one possible neural mechanism for these syllable transitions. In this model, transitions are mediated by neural connections within HVC that extend from the tail of one chain to the heads of subsequent chains (Jin, 2009). Each state in a POMM corresponds to a single syllable chain. Hence the multiple states for a syllable correspond to multiple syllable chains that drive the production of the same syllable. For example, consider Example 1, illustrated in Fig.1, where there are two unique observed sequences, *ACD* and *BCE*. The POMM designates two states for syllable *C*, requiring two distinct syllable-chains for *C*, as depicted in Fig.6a.

The fact that deafening reduces state multiplicity rules out the intrinsic mechanism in which syllable transitions do not rely on auditory feedback. An alternative is the reafferent mechanism (Sakata and Brainard, 2006; Sakata and Brainard, 2008; Hanuschkin et al., 2011; Wittenbach et al., 2015). Here, auditory feedback from preceding syllables strongly influences syllable transitions. This allows for the possibility that all syllables are produced by single syllable-chains, regardless of state multiplicity.

We illustrate the reafferent mechanism with Example 1 (Fig. 6b). In this case, there is one syllable-chain for *C*, which connects to chain-D and chain-E. However, the activations of chain-D and chain-E are determined by the reafferent auditory inputs (Sakata and Brainard, 2006; Sakata and Brainard, 2008; Hanuschkin et al., 2011; Wittenbach et al., 2015). The auditory feedback from syllable *A* is sent to chain-D, while the auditory feedback from syllable *B* is sent to chain-E (Fig. 6b). The auditory inputs can bias the transitions to chain-D or chain-E from chain-C (Jin, 2009; Hanuschkin et al., 2011; Wittenbach et al., 2015). With strong enough auditory inputs, the probability of transition from *C* to *D* should approach 1 when *C* is preceded by *A*. Conversely, when *C* is preceded by *B*, the transition probability to *E* should approach 1.

Two pieces of evidence cast doubt on the reafferent mechanism. First, while deafening significantly reduces the state multiplicity in the POMMs, it does not completely abolish the state multiplicity, suggesting that multiple states cannot be entirely due to auditory feedback. Second, the durations of syllables and the silent gaps between syllables can be longer than the time it takes for auditory feedback to reach HVC, such that by the time of the transitions to the next syllables, the auditory feedback from preceding syllables has faded away and thus is incapable of providing context-dependent syllable transitions.

We illustrate the second point with the example in Fig. 6b. For the activity in chain-A to bias the transition from chain-C towards chain-D, the feedback signal from syllable *A* must be available at the time of the transition. Thus, the delay of the “round-trip” from the activity in chain-A to the auditory feedback in HVC from syllable A must be longer than the duration of chain-C. The duration of the activity propagation in chain-C should correspond to the duration of syllable *C* and the preceding silent gap.

In Fig. 7, we plot the distributions of the gap-syllable durations for all syllables that require multiple states under the normal condition for the birds. The median values of these durations approximately range from 80 ms to 180 ms. Some syllables are repeated; in these instances, the auditory feedback must be available after the completion of the syllable repetitions. The distributions of the durations of gap-syllables, including repetitions, are shown in Supplementary Fig. S6, where the median values of the durations extend approximately from 80 ms to 600 ms. These ranges establish a lower limit on the delays of the auditory feedback necessary for the reafference mechanism to function effectively.

Experiments that perturbed auditory feedback during singing in Bengalese finches found that perturbation of sound altered HVC activity approximately 44 ms after the onset of the perturbation, with no significant changes in HVC activity detectable after 80 ms (Sakata and Brainard, 2008). Given that the premotor delay from HVC to syllable production is around 50 ms (Schmidt, 2003), the round-trip delay of auditory feedback from a syllable should be limited to 130 ms. Therefore, the reafference mechanism can be ruled out for at least 28% of syllables with state multiplicity, as the median durations of gap-syllables for these syllables exceed 130 ms (Fig. 7). This fraction increases to 44% when syllable repetitions are accounted for (Supplementary Fig. S6).

An alternative mechanism does not necessitate long delays of auditory feedback. In this mechanism, multiple states are encoded by separate chains, similar to the intrinsic mechanism, but transitions between the chains are influenced by auditory feedback. If auditory feedback “tunes” the transition probabilities between the chains to create context dependencies in syllable transitions, then deafening will “de-tune” them, reducing the context dependencies. In Fig. 6c, we illustrate this auditory-tuning mechanism with Example 1. The auditory feedback from chain-A biases the transition to the chain-C that transitions to chain-D, whereas the auditory feedback from chain-B biases the transition to the chain-C that transitions to chain-E. Loss of auditory feedback causes the network to revert to relying solely on the intrinsic connections, which permits transitions from chain-A and chain-B to both chain-Cs.

To investigate this possibility, we consider each state in the POMMs under the normal condition as syllable-chains, then modify the transition probabilities between them. These modifications are informed by the pairwise syllable transition probabilities observed after deafening. If a novel transition from syllable *x* to syllable *y* emerges after deafening (with a probability of *≥* 0.01), new transitions are established from all states associated with *x* to all states associated with *y*. Conversely, if the transition from *x* to *y* is eliminated after deafening, we remove the transitions from all states associated with *x* to all states associated with *y*. We further refine the transition probabilities between the states. From a state associated with *x*, if transitions exist to states associated with *y*, the probabilities of these transitions are scaled to ensure the total transition probability matches the transition probability from *x* to *y* after deafening. These modifications of the transition probabilities could diminish the context dependencies, potentially simplifying the POMMs to versions with reduced state multiplicity (referred to as reduced POMMs). It is important to note that these modifications do not increase state multiplicity, as the number of syllable-chains remains constant and no new states are introduced.

The reduction process unfolds as follows: With a modified POMM, we generate 100 sets of *N* sequences, where *N* represents the number of sequences observed for the deafened bird. Among these, we select the set with the maximum probability given the modified POMM as the “observed syllable sequences”. Using these “observed sequences”, we infer minimal POMM, which represent the reduced POMM after modifying the transitions. This process is illustrated in Fig. 8a-c, using o10bk90 as an example. The reduced POMM for this bird matches the POMM after deafening in terms of the number of states for each syllable, although the transition probabilities differ in detail due to the fluctuations introduced by the finite *N*. Similarly, the reduced POMMs for all other birds align with the POMMs post-deafening, with the exception of o46bk78, where syllable *c* has an additional state, as shown in Supplementary Figs. S7-S8.

**Figure 6.**
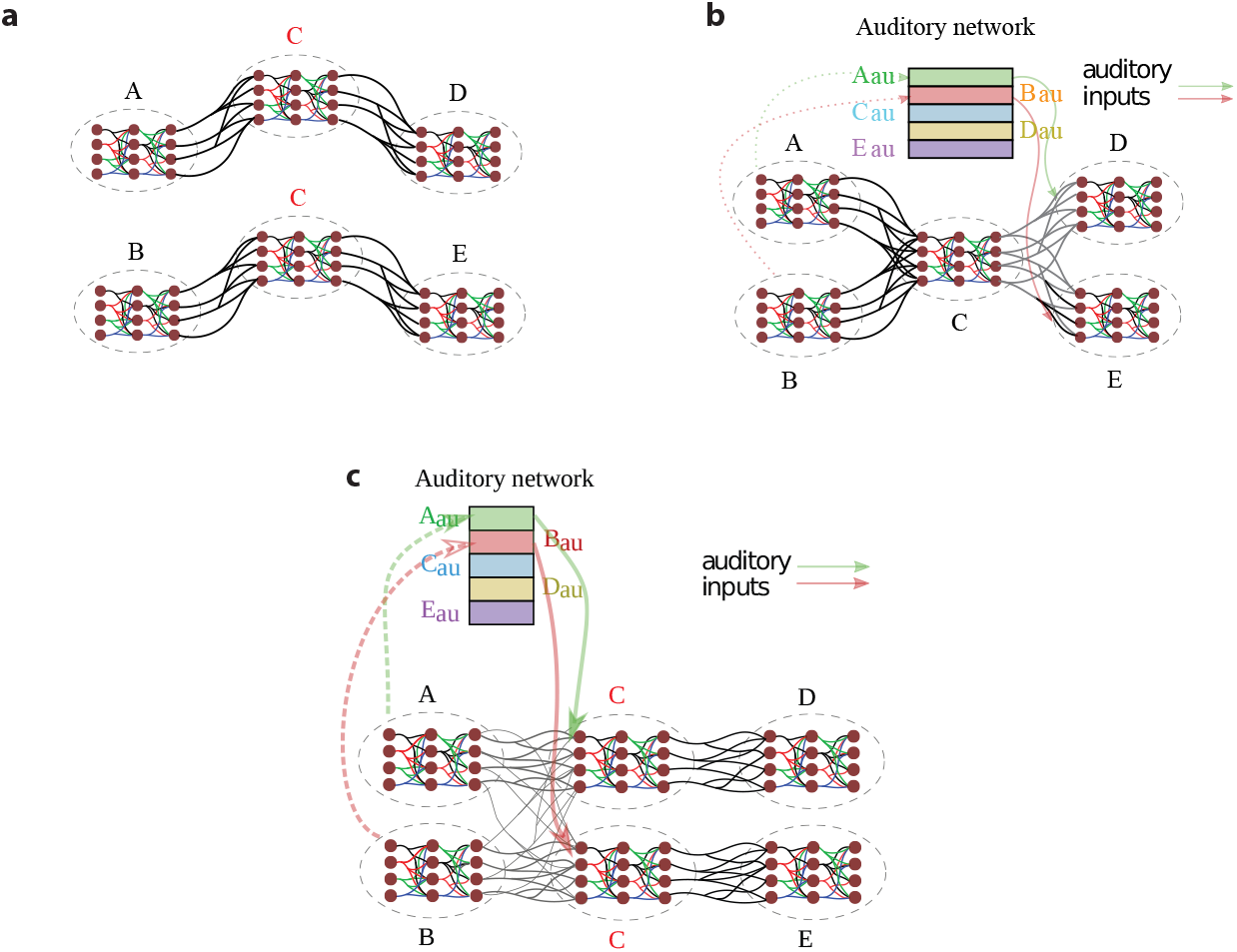
Neural mechanisms for the POMM in Example 1 (Fig. 1). There are two unique sequences, *ACD* and *BCE*, in the observed set. **a**. The intrinsic mechanism for the multiple states of syllable *C*. Two syllable-chains encode the two states for *C*. **b**. The reafference mechanism for the multiple states of syllable *C*. There is one syllable-chain for *C*, with the multiple states arising due to the auditory feedback from preceding syllables differentially influencing the syllable transitions from *C*. **c**. The auditory-tuning model for the multiple states of syllable *C*. There are two syllable-chains for *C*. Intrinsic connections exist from chain-A and chain-B to the two chain-Cs. Auditory feedback from syllable A biases the transition from chain-A to the upper chain-C, while auditory feedback from syllable B biases the transition from chain-B to the lower chain-C.

**Figure 7.**
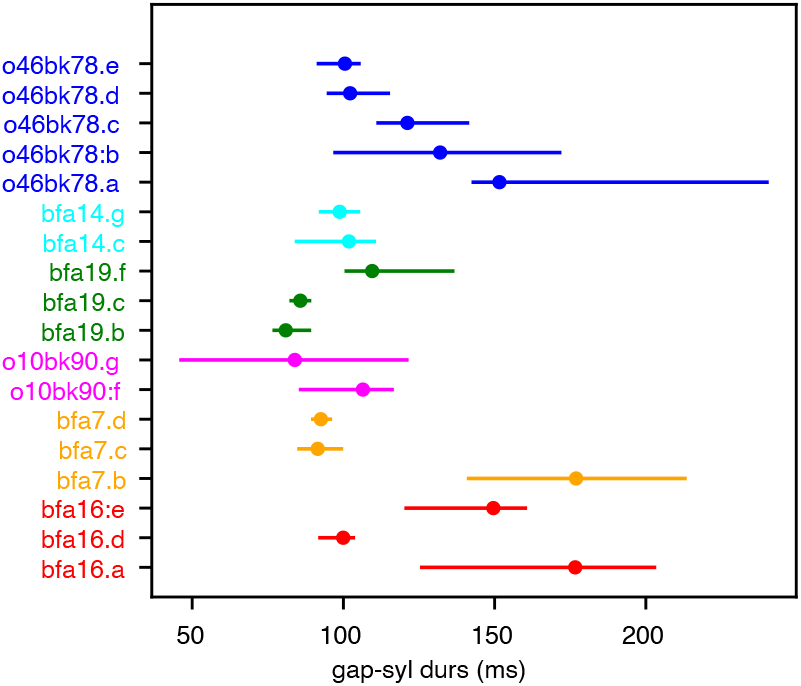
Distributions of gap-syllable durations. Durations of syllables plus the preceding gaps are shown for all syllables with multiple states in the POMMs for all birds. The dots indicate the median values and the bars indicate the 5% - 95% ranges of the distributions.

**Figure 8.**
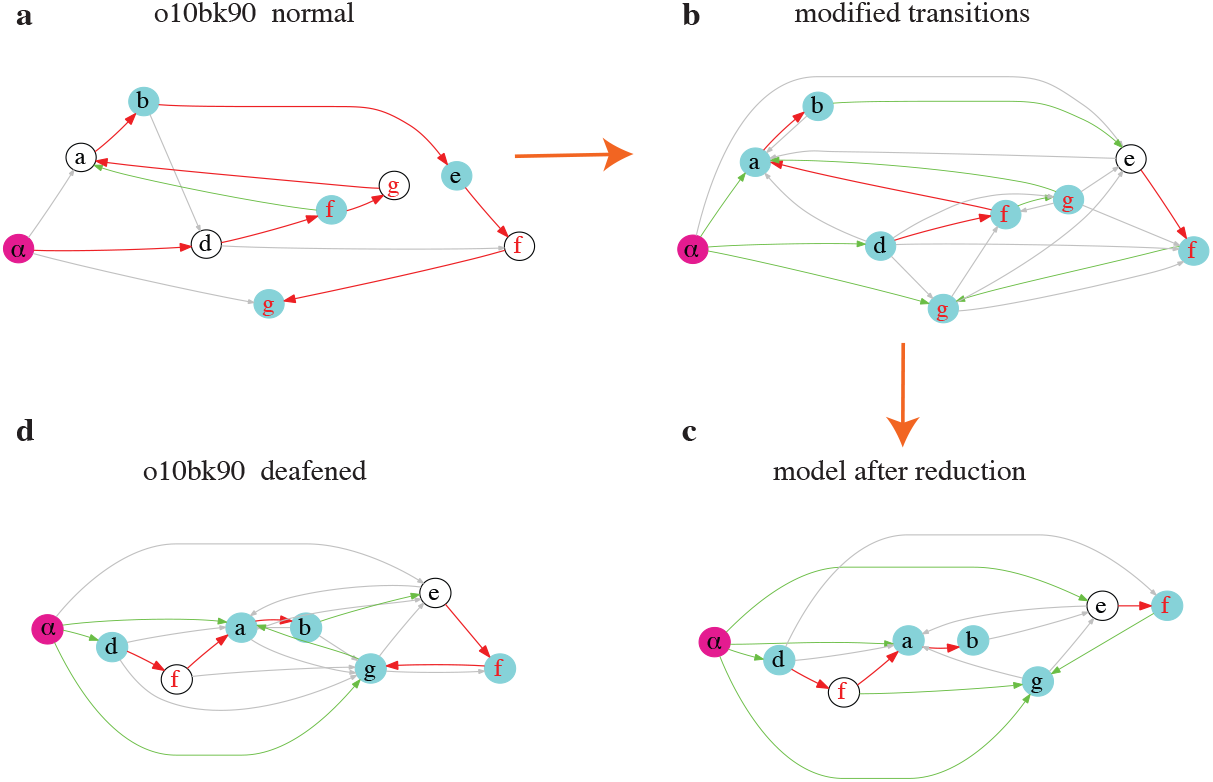
Reduction of a POMM after modifications of the transition probabilities. **a**. The POMM of o10bk90 in the normal condition. **b**. The POMM after modifications of the transitions probabilities using the syllable transition probabilities after deafening. **c**. Reduced POMM inferred from the *N* = 62 “observed sequences” generated from modified POMM. **d**. The POMM after deafening shown for comparison with the reduced POMM.

For the three birds whose POMMs become Markovian after deafening (bfa7, bfa19, bfa14), merely adding the novel transitions and removing the deleted ones – without further modifying the transition probabilities between the states beyond the necessary normalizations to ensure that the total transition probability from a state equals 1 – is sufficient to render the reduced POMMs Markovian (Supplementary Fig. S9). For o46bk78, the number of extra states is also significantly reduced. However, for the other two birds (bfa16, o10bk90), this approach does not alter the number of extra states in comparison to the POMMs under the normal condition.

## Discussion

Like human language, birdsong is characterized by context-dependent transitions in syllable sequences. POMMs are generative models suitable for birdsong syllable sequences (Jin and Kozhevnikov, 2011). POMMs capture context dependencies by associating multiple states with the same syllable. From the start state to the end state, a state transition path generates a syllable sequence. A syllable with state multiplicity can transition differently depending on the state it is in, thus being involved in context-dependent transitions to other syllables. POMMs lacking sufficient state multiplicity may tend to overgeneralize, creating sequences that are never observed (type I context dependence) or may increase probabilities of certain sequences more than observed (type II context dependence). By statistically testing whether a POMM overgeneralizes compared to the observed set of syllable sequences, we can infer a minimal POMM that has the smallest number of states among POMMs that are compatible with the observed set. By analyzing the minimal POMMs for Bengalese finch songs under the normal condition and shortly after deafening, we show that auditory feedback plays an important role in creating context-dependent syllable transitions in Bengalese finch songs.

To evaluate the fit of a POMM, we construct the distribution of the augmented sequence completeness *P*_*β*_ for the sequences sampled from the candidate POMM. This distribution is used to calculate the *p*-value of the *P*_*β*_ of the observed set computed with the POMM. We use the criterion *p <* 0.05 to reject the POMM. Making the rejection criterion more stringent by lowering this *p*-value cut-off should allow for the acceptance of POMMs with fewer extra states.

Our method is computationally intensive, primarily due to the significant computational effort required in determining the minimal number of states for the syllables. Investigating more efficient methods for estimating state multiplicity presents a compelling avenue for future research. A promising approach involves quantifying the predictive information within syllable sequences to infer the requisite number of parameters for accurately encoding sequence complexity (Bialek et al., 2001).

Our method is conservative, especially when the number of observed sequences *N* is small. In such cases, POMM tends to underestimate the true number of states due to the insufficient representation of context dependencies in the observed sequences. A practical approach to determine if *N* is sufficiently large involves examining whether the sequence completeness *P*_*c*_, as calculated using POMM, is near 1 for the observed sequences. The difference 1 *− P*_*c*_ can then provide a rough estimate of the probability associated with missing unique sequences.

In our earlier work, POMMs were derived by aligning them with sequence analysis metrics such as n-gram distributions, which represent the probabilities of n-length subsequences (Jin and Kozhevnikov, 2011). This prior method often resulted in an overestimation of the necessary state multiplicity, a tendency exacerbated by small sample size *N*. Additionally, simplifying the model frequently necessitated manual adjustments. Our current approach marks a significant departure from these constraints, demonstrating robustness even when *N* is small. This improvement stems from the method’s reliance on sequence completeness – a cumulative probability metric – which is insensitive to the precision of individual probability estimates. Despite not directly conforming to n-gram distribution fitting, our method ensures that the statistical properties of 2-to 7-gram sequences generated by the POMM align with those observed in the sequences of all birds studied (Supplementary Fig. S10 and S11).

Two common methods of model selection are the Akaike Information Criterion (AIC) and the Bayesian Information Criterion (BIC) (Zucchini and MacDonald, 2009). For a POMM, the AIC is defined as 2*k −* 2*L*, where *k* is the number of transition probabilities, and *L* is the log-likelihood of the observed sequences given the POMM. The BIC is defined as *k* log *N −*2*L*, where *N* is the number of observed sequences. Among candidate POMMs, the one with the minimal AIC or BIC is selected to balance model fit and complexity. When tested with a simple example with a known ground truth model, as shown in Fig. 2, we find that AIC and BIC underestimate the state multiplicity compared to our method (Supplementary Fig. S12). AIC and BIC must compare between models and often require enumerating possible models to select the minimal model. In contrast, our method can evaluate a single model directly. This approach makes it possible to initially accept a complex model and then simplify it through merging and deleting states. Since our method does not use gradient information, it is not hampered by the issue of local minima in optimization approaches.

In the normal condition, the songs of Bengalese finches in our study exhibit context-dependent syllable transitions. The POMMs for these songs demonstrate varying levels of state multiplicity, as illustrated in Figs. 3-4, indicating significant individual differences. Investigating the origins of these differences could be interesting, particularly in determining the extent to which learning influences the formation of context dependencies in the songs.

Deafening significantly reduces the state multiplicity, yet, notably, it does persist for certain syllables in some birds. This observation indicates that context-dependent syllable transitions are not solely reliant on auditory inputs. It suggests that an intrinsic mechanism, as illustrated in Fig. 6a, contributes at least partially to the formation of context dependencies.

The observed reduction in state multiplicity following deafening appears to support the reafference mechanism (Fig. 6b). Yet, the applicability of this mechanism is circumscribed by the duration for which auditory feedback from the context-providing syllables is retained within the HVC. Research involving the disruption of auditory feedback in Bengalese finches has demonstrated that such perturbations modulate HVC activity, with the impact manifesting after delays, ceasing around 80 ms after the perturbation (Sakata and Brainard, 2008). Considering an approximate premotor delay of 50 ms from HVC activity to the actual production of a syllable (Schmidt, 2003), it follows that the auditory feedback from a context-providing syllable would be accessible for a maximum of 130 ms, beyond which it would not. This temporal boundary narrows the “distance” over which context dependence can operate, essentially the span of gap+syllables that can be interposed between context-providing syllables and those undergoing context-dependent transitions. Analysis of durations of gap+syllables indicates a maximum feasible distance of one syllable, with numerous instances where even one gap+syllable duration exceeds the temporal limit, as evidenced in Fig. 7 and Supplementary Fig. S6. Consequently, the finite window for auditory feedback imposes a considerable constraint on the effectiveness of the reafference mechanism in facilitating context-dependent syllable transitions.

The auditory-tuning mechanism incorporates aspects of both the intrinsic and reafference mechanisms, as depicted in Fig. 6c. Similar to the intrinsic mechanism, a syllable represented by *m* states in a POMM corresponds to at least *m* syllable-chains within the HVC. Echoing the principles of the reafference mechanism, auditory feedback modulates transitions between syllable-chains, specifically targeting those transitions that directly follow the context-providing syllables. While intrinsic connections among syllable-chains exist, they alone do not account for the majority of context dependencies observed in syllable sequences. Instead, auditory feedback refines the transition probabilities, enhancing context dependencies by diminishing the randomness in transitions prompted by intrinsic connections. This effect is underscored by observations that disruptions to auditory feedback during song in Bengalese finches generally increase the randomness of syllable sequences and can introduce novel transitions (Sakata and Brainard, 2006). Consistent with the auditory-tuning mechanism, adjusting the POMMs under the normal condition to reflect syllable transition probabilities measured post-deafening significantly aligns these POMMs with those observed following deafening (Fig. 8 and Supplementary Figs. S7-S8).

Calcium imaging techniques in the HVC of singing birds have revealed the many-to-one mapping from syllable-chains to syllables (Cohen et al., 2020). When applied to the Bengalese finch, this approach should enable a quantitative assessment of POMMs within the framework of the auditory-tuning mechanism. For each Bengalese finch, POMMs can be derived from their unique syllable sequences. The identified number of states for each syllable within these models serves as a minimal estimate for the corresponding number of syllable-chains detectable in the HVC through calcium imaging. This method provides a direct means to test the accuracy of POMMs by comparing the theoretical predictions with empirical data obtained from individual birds.

Previous studies on the effects of deafening in the Bengalese finch have highlighted the rapid loss of sequence stereotypy following deafening, underscoring the necessity of online auditory feedback for maintaining stereotyped syllable sequences (Woolley and Rubel, 1997; Okanoya and Yamaguchi, 1997). Our findings are consistent with these observations, as we also observe that syllable sequences become more randomized post-deafening. On average, across the birds studied, there is a noticeable increase in transition entropy at the syllable transition branching points after deafening (Fig. 5b). This increase is primarily attributed to the previously dominant transitions at these branching points becoming more evenly distributed, leading to branches with more similar transition probabilities. Notably, similar outcomes have been documented in studies involving perturbations of auditory feedback (Sakata and Brainard, 2006), cooling of the HVC (Zhang et al., 2017), and the enhancement of inhibition within the HVC (Isola et al., 2020) in Bengalese finches. Exploring the potential for a unified neural mechanism underlying these diverse manipulations presents an intriguing avenue for further research.

In conclusion, we have developed a method for inferring minimal POMMs based on observed sequences. Applying this method to the syllable sequences of Bengalese finch songs, both before and after deafening, indicates that the auditory system plays a significant role in establishing context dependencies in syllable transitions. This approach should be widely applicable to the analysis of behavioral sequences in other animals.

## Materials and Methods

The data set in this work was previously used for analyzing syllable repeats in Bengalese finch songs (Wittenbach et al., 2015) (download from http://www.dezhejinlab.org/SharedData/). Details of recording songs, annotating syllables, and deafening through bilateral cochlear removal, as well as the Ethics Statement can be found in the published paper (Wittenbach et al., 2015). We specifically used the data collected from six male adult Bengalese finches before and after deafening (labeled bfa14, bfa16, bfa19,bfa7, o10bk90, and o46bk78). For comparing distributions of paired data in Fig. 5, we use Wilcoxon signed-rank test using scipy.stats.wilcoxon, which is in the Python module scipy (Virtanen et al., 2020). Mathematical details of generating sequences from POMMs and inferring minimal POMMs are in Supporting Information.

## Acknowledgements

Research was supported by NSF award EF-1822476 (DZJ). The funders had no role in study design, data collection and analysis, decision to publish, or preparation of the manuscript

## Supporting Information

### Data set

In the data set, syllables are labelled *a* through *l*, and *x* through *z*. Some ambiguous syllables are noted with symbols 0 and *−*, and they are skipped. Bengalese finch song bouts typically begin with short introductory notes. They are labeled as *i, j* and *k*. We define song sequences as segments of syllables that are bracketed by periods of introductory notes and the end of the recordings.

### POMM

A POMM is specified by a state vector *V* = [*α, ω, s*_3_, *s*_4_, *· · ·, s*_*n*_], where *s*_1_ = *α* and *s*_2_ = *ω* are the start and the end states, *n* is the total number of states, and *s*_*i*_ for *i* = 3, *· · ·, n* is the syllable symbol associated with the *i*th state. The same syllable symbol can appear multiple times in the state vector. Transitions between the states are described by a transition matrix *T*, whose element *T*_*ij*_ gives the probability of transition from state *i* to state *j*. There are no transitions to the start state, i.e. *T*_*i*1_ = 0; and there are no transitions from the end state, i.e. *T*_2*j*_ = 0.

Sequence generation from a POMM starts with the start state. At state *i*, the next state *j* is chosen with the probabilities *T*_*ij*_ among possible choices of state 2 to state *n*. Once chosen, the symbol *s*_*j*_ is added to the sequence. This process repeats until the end state is reached, at which point the sequence generation is complete.

A POMM is visualized with the software Graphviz (Ellson et al., 2001). To reduce clutter, only transitions with probabilities larger than 0.01 are shown. Additionally, the End-state is not shown. Instead, the states that can transition to the end state are shown in cyan. The transition probability from one state to the end state is 1 minus the sum of the transition probabilities to other states. If a state does not transition to the end state with a probability larger than 0.01, the state is shown as white. The start state is shown in pink.

### Markov model

A Markov model is a special case of POMM for which each syllable symbol appears only once in the state vector. The transition probabilities *T* can be computed as

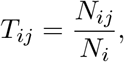

where *N*_*i*_ is the total number of times *s*_*i*_ appears in the set *Y* of sequences, and *N*_*ij*_ is the total number of the times that the two-symbol subsequence *s*_*i*_*s*_*j*_ appears in *Y*. Note that

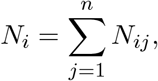

so we only need to compute *N*_*ij*_.

### Baum-Welch algorithm

Computing *T* for POMM with state multiplicity is more complicated than that for the Markov model, but the approach is similar. Starting from a set of random transition probabilities, the state transition sequences that correspond to the syllable sequences in *Y* are worked out. The transition probabilities are then updated according to

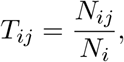

where *N*_*i*_ is number of times the state *i* appears in the state sequences, and *N*_*ij*_ is the number of times the subsequence of states *ij* appears. With the updated *T*, the process is repeated. The process stops when the changes in *T* is smaller than 10^*−*6^. Because the result might be dependent on the initialization of *T*, the process is run for 100 times with different seeds for random number generator. The *T* that maximizes the probabilities of generating *Y* from the POMM is selected.

The computation is efficiently implemented with the Baum-Welch algorithm (Rabiner, 1989). Consider a sequence *y*_1_*y*_2_ *· · · y*_*t*_ *· · · y*_*m*_ in the set *Y*. Here *t* is the step in the sequence and *m* is the maximum length of the sequence. The algorithm consists of three parts. First, calculate the forward probability *α*_*i*_(*t*), which is the probability of being at state *i* at step *t* given the proceeding sequence is *y*_1_*y*_2_ *· · · y*_*t−*1_. This is computed iteratively with

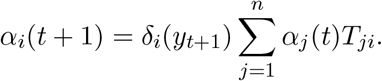

Since all sequences start from the Start-state, the initial condition is *α*_1_(0) = 1 and *α*_*j*_(0) = 0 for all *j ≠* 1. Here *δ*_*i*_(*y*_*t*+1_) = 1 if the symbol *y*_*t*+1_ at step *t*+1 is the same as the symbol *s*_*i*_ associated with state *i*; otherwise, *δ*_*i*_(*y*_*t*+1_) = 0. Second, calculate the backward probability *β*_*i*_(*t*), which is the probability being at state *i* at step *t* and the follow-up sequence is *y*_*t*+1_, *· · ·, y*_*m*_. This is calculated iteratively with

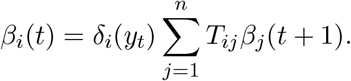

Since all sequences end at the end state, the initial condition is *β*_2_(*m* + 1) = 1 and *β*_*j*_(*m* + 1) = 0 for all *j ≠* 2. Third, calculate *N*_*i*_ and *N*_*ij*_. The forward and backward probabilities *α*_*i*_(*t*) and *β*_*i*_(*t*) should be computed for each sequence in *Y*. The number of transition from state *i* to state *j* is given by

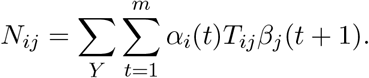

For a given sequence *y*_1_*y*_2_ *· · · y*_*m*_, the probability that the POMM generates it is given by

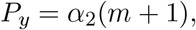

which is the forward probability of ending at the end state at step *m* + 1.

The total probability of the set *Y* is given by

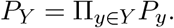

It is most convenient to use the log-likelihood, which is

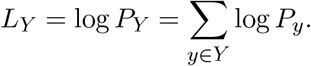

### Sequence completeness, total variation distance and augmented sequence completeness

For a set of sequences *Y*, the sequence completeness on a POMM is computed as

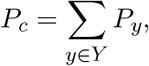

where *y* is a unique sequence in *Y*. The sum is over all the unique sequences in the set.

For a set of observed sequences *Y*_*o*_, the total variation distance is defined as

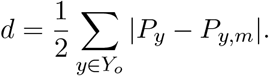

Here *P*_*y,m*_ is the probability of the unique sequence *y* computed on the POMM and then *normalized* among the unique sequences such that

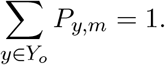

This normalization is necessary because *P*_*y,m*_ is compared to *P*_*y*_, which is normalized:

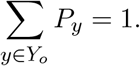

The total variation distance ranges from 0 to 1.

The augmented sequence completeness is defined as

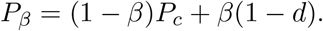

Here *β* is a parameter that can be chosen in the range (0, 1). The value of *P*_*β*_ ranges from 0 to 1. A perfect POMM for the observed set should yield *P*_*β*_ = 1 because *P*_*c*_ = 1 and *d* = 0. When *N* is small, the measurements of *P*_*y*_ are not accurate. For this case, the contribution from *d* should be reduced by taking a small value for *β*. In our work, we chose *β* = 0.2.

### Statistical test

To test whether an observed set *Y*_*o*_ with *N* sequences could be generated from a POMM, we generate *M* = 10000 sets of *N* sequences, and compute the *P*_*β*_ of the generated sets, which gives a distribution of *P*_*β*_. We also compute the augmented sequence completeness *P*_*β,o*_ of the observed set. In the distribution, we count the number *K* of *P*_*β*_ that are smaller or equal to *P*_*c,o*_. To avoid small fluctuations in *P*_*β*_ making *K* artificially small, we added 10^*−*10^ to *P*_*β,o*_. The p-value is

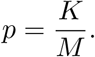

The POMM is rejected if *p <* 0.05, and accepted otherwise.

### Inferring minimal POMM

For a given set *Y* of *N* syllable sequences, the minimal POMM is inferred through three steps: setting up the initial POMM through higher order Markov models; simplifications with state merging followed by state deletion; and removal of transitions. Because of the inherent randomness in the inference process, the minimal POMMs from different runs may differ. We run the process 20 times and select the most common model as the final minimal POMM.

The initial POMM is constructed to be equivalent to a higher order Markov model for *Y*. We flank each sequence with the start symbol *α* and the end symbol *ω*, and find all subsequences of length *m*. Here *m* is the order of the Markov model. We also include all subsequences starting with the symbol *α* and the lengths from 1 to *m −* 1. We identify all unique subsequences. Each unique subsequence is assigned a state, and the last symbol of the subsequence is associated with the state. We count the number of times *N*_*i*_ that the *i*th subsequence appears, as well as the number of times *N*_*ij*_ that the *i*th subsequence is followed by the *j*th subsequence. The transition probability from state *i* to state *j* is set to

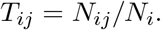

This completes the construction of the POMM that is equivalent to the *m*th order Markov model. Starting from *m* = 1, we increase *m* until the POMM passes the statistical test. This completes the initialization of the POMM.

To simplify the POMM, we first test merging states and then test deleting states. Merging states *i* and *j* with the same associated syllable eliminates state *j*. The transition probability from state *i* to state *k* (*k ≠ i, j*) is recomputed as

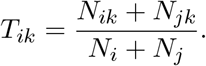

The transition probability from state *k* (*k ≠i, j*) to state *i* is updated as

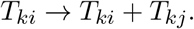

The number of times state *i* is visited is updated as

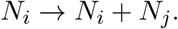

If the POMM after the merger passes the statistical test, the merger is accepted. Otherwise the merger is reversed. To speed up this process, we first test merging all states for each syllable. If the merger is rejected, we test pairwise merging of the states for the syllable. After no further state mergers are possible, we test deleting states. For each syllable, we reduce the number of states by 1. We then compute the transition probability matrix *T* using the Baum-Welch algorithm to maximize the log-likelihood of *Y*. If the POMM passes the statistical test, the reduction continues. Otherwise the reduction is reversed. The process continues until no further state deletions are accepted.

The final step is minimization of the number of transitions in the POMM. We first remove all transitions with probability smaller than 0.001. We then remove the remaining transitions one by one, and re-compute the transition matrix *T* after each removal. To remove a transition from state *i* to state *j*, we set *T*_*ij*_ = 0 in the initial transition matrix for the Baum-Welch algorithm. The algorithm ensures that this transition element remains 0. If the log-likelihood remains within the threshold, the removal is accepted; otherwise the removal is reversed. The threshold is

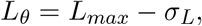

where *L*_*max*_ is the log-likelihood of the original POMM before any removals, and *σ*_*L*_ is the standard deviation of the log-likelihood of the 100 runs of Baum-Welch algorithms with different random seeds. If after all of the accepted removals the POMM does not pass the statistical test, we revert to the POMM before any removals.

### Probability of finding a subsequence

The probability *P*_*s*_ of finding a subsequence in a set *Y* is defined as

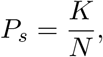

where *N* is the number of sequences in the set, and *K* is number of sequences that contains the subsequence.

### State merging tests

To evaluate the context-dependent syllable transitions encoded by state multiplicity in a POMM, we perform pairwise state merging tests. The merged state retains all transitions to and from the two states. The transition probabilities of the state-merged POMM are recomputed using the Baum-Welch algorithm and the observed set *Y*_*o*_. By examining the states transitioning into the two states, and the states that follow the two states, we find possible subsequences that can show overgeneralization after the state merger. We find a subsequence that either is unseen in the observed set (*P*_*s,o*_ = 0) or has small probability *P*_*s,o*_. To see whether the subsequence is significantly more probable in the sequences generated from the state-merged POMM, we generate 10000 sets of *N* sequences from the POMM. Here *N* is the number of sequences in *Y*_*o*_. For each generated set, we compute *P*_*s*_. This creates a distribution. We count the number of *P*_*s*_ that is smaller than or equal to *P*_*s,o*_ + 10^*−*10^. The p-value is the ratio of this number and 10000. We add a small number 10^*−*10^ to *P*_*s,o*_. This is for avoiding artificially lowering p-value due to those *P*_*s*_ that are equal to *P*_*s,o*_. For example, if the subsequence is unobserved (*P*_*s,o*_ = 0) and the state-merged POMM does not generate it either, we would have a situation that *P*_*s*_ = 0 for all of the sampled set. By adding the small number to *P*_*s,o*_, we ensure that *p* = 1, as it should be. If *p <* 0.05, we conclude that the enhancement of *P*_*s*_ after state merger is significant.

### Analysis of context dependent syllable transitions

In the following, we show such dependencies for each bird in the normal condition and after deafening. We first show the major syllable transitions in the observed sets. We then point out how reducing the state multiplicity by merging states associated with the same syllable makes the POMM overgeneralize or significantly enhance the probabilities of some subsequences compared to the observed. This merging technique is inspired by the examples shown in Fig. 1. The state-merged POMM retains all state transition branches of the original POMM, but the transition probabilities are re-calculated with the Baum-Welch algorithm using the sequences in the observed sets. Since our goal is to show why multiple states are needed in the original POMMs, we select a few subsequences for evaluation and do not discuss all possible context dependencies in the syllable sequences.

The evaluation is done with the measure *P*_*s*_, which is the fraction of sequences in a set that contain a selected subsequence. We first compute *P*_*s*_ for the set of *N* observed sequences. We then compute *P*_*s*_ for 10000 sets of *N* sequences generated from a state-merged POMM. This creates a distribution of *P*_*s*_ from the generated sets. We report the median *P*_*s*_ of this distribution to show how much the probability is enhanced compared to the *P*_*s*_ for the observed set. The significance of the enhancement is shown with the p-value, which is the probability *p* that *P*_*s*_ in the distribution is smaller than the *P*_*s*_ for the observed set. The process is analogous to the test of POMMs shown in Fig. 2.

### bfa7

In the normal condition, syllable *b* has 2 states and syllables *c* and *d* have 4 states each (Fig. 3). The two states for *b* encode the following context dependence:

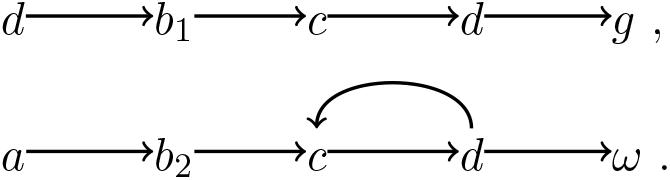

The state multiplicity for *c* and *d* reflects the following context dependencies:

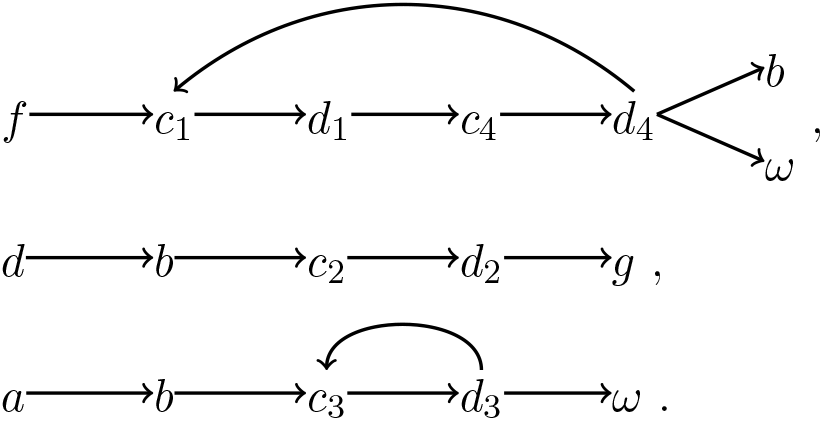

Here *ω* denotes the end of the sequence, and the subscripts indicate different states for the same syllable. The consequences of merging states are summarized in Table S1 (Supporting Information). The mergers make the POMM overgeneralize, generating subsequences that are unobserved (for example, *f → c → d → g*). These are examples of type I context dependencies.

Deafening leads to the appearance of *c → a* and *h → a* transitions, strengthening of *d → a* transition, and disappearance of *d → c* transition. Except for *b*, the sequence can now stop at all syllables. Interestingly, the *d → g* transition is lost and syllable *g* does not appear after deafening. The syntax is Markovian, suggesting that there is no context dependence.

### bfa14

In the normal condition, the POMM has two states for *c* and *g* (Fig. 3). The two state for *c* reflect the following type II context-dependent syllable transitions:

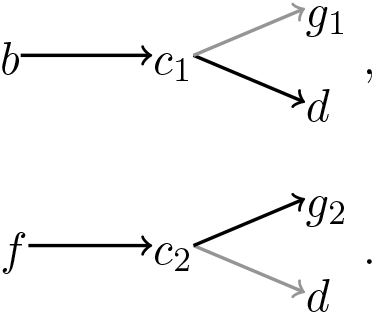

Transition from *c*_1_ to *d* is strong (transition probability *P* = 0.9), but to *g* is weak (*P* = 0.08). On the contrary, transition from *c*_2_ to *d* is weak (*P* = 0.05), but to *g* is strong (*P* = 0.9). Merging *c*_1_ and *c*_2_ significantly enhances the probability of the subsequence *b → c → g* from *P*_*s*_ = 0.07 (observed) to *P*_*s*_ = 0.2 (*p* = 0); and *f → c → d* from *P*_*s*_ = 0.01 (observed) to *P*_*s*_ = 0.1 (*p* = 0).

Transition from *g*_1_ to *e* is stereotypical (*P* = 1.0). In contrast, transition from *g*_2_ to *e* is weak (*P* = 0.03); instead, sequence is most likely stop at *g*_2_ (*P* = 0.97 for *g*_2_ *→ ω*). Merging *g*_1_ and *g*_2_ enhances the probability of the subsequence *f → c → g → e* from *P*_*s*_ = 0.01 (observed) to *P*_*s*_ = 0.05, which is significant with *p* = 0.03.

Deafening creates numerous novel transitions with small probabilities (*<* 0.1) (Fig. 3). Novel transitions with large probability (*≥* 0.1) also occur, which include transitions *a → h, b → l, h → f*, and *l → g*, as well as from *α* to syllables *b, c, e, f, l*. Some transitions are weakened, which include transitions *l → c* and *f → c*. All context dependencies are lost, and the model becomes Markovian.

### bfa16

In the normal condition, there are two states for syllables *a, d*, and *e* (Fig. 3). The two states for *a* encode the following context dependence:

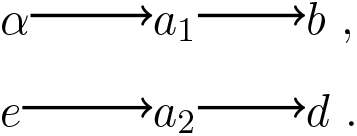

Here *α* denotes the start of the sequence. Merging *a*_1_ and *a*_2_ creates an unobserved subsequence *α → a → d* with median *P*_*s*_ = 0.2 and *p* = 0.

The two states for *d* encodes the following context dependence:

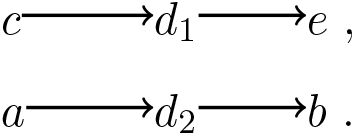

Merging *d*_1_ and *d*_2_ creates an unobserved subsequence *c → d → b* with median *P*_*s*_ = 0.2 and *p* = 0.

The two states for *e* encodes the context dependence

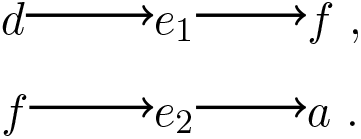

Merging *e*_1_ and *e*_2_ creates unobserved subsequence *d → e → a* with median *P*_*s*_ = 0.4 and *p* = 0.

The major effects of deafening are the loss of the transition from *e*_2_ to *a*_2_; the strengthening of the transition *e*_2_ *→ a*_1_; and the enhancement of stopping after *g*. The only state multiplicity left is for syllable *e*, which encodes the same context dependency as in the normal condition. Merging the two states for *e* again creates unobserved subsequence *d → e → a* with median *P*_*s*_ = 0.04 and *p* = 0.

### bfa19

In the normal condition, there are two states for syllables *b, c*, and *f* (Fig. 4). The state multiplicity for *b* and *c* encodes the context dependence

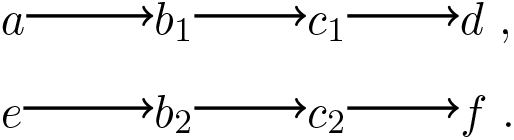

Merging either *b*_1_ and *b*_2_ or *c*_1_ and *c*_2_ creates unobserved subsequence *a → b → c → f* with median *P*_*s*_ = 0.4 and *p* = 0.0001.

The state multiplicity for *f* encodes the context dependence

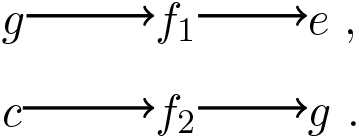

Merging *f*_1_ and *f*_2_ creates unobserved subsequence *g → f → g* with median *P*_*s*_ = 0.3 and *p* = 0.0009.

After deafening, many novel transitions appear, most notably *a → d*. Sequences can also start with *d* or *e* with probabilities greater than 0.1. The model becomes Markovian, and all context dependencies disappear.

### o46bk78

In the normal condition, the POMM has multiplicity for multiple syllables (Fig. 4). There are 4 states for *b*, 3 states for *a* and *c*, and 2 states for *d* and *e*, respectively. The structure of the POMM can be summarized as two motifs connected with a bridge. Motif 1 is the following cycle:

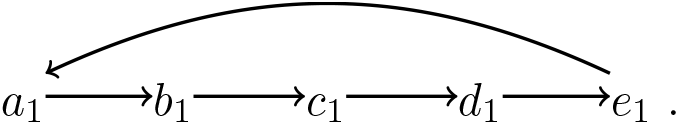

The start state transitions to *a*_1_. The bridge is the subsequence

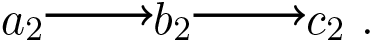

Motif 1 transitions to the bridge via the transitions *b*_1_ *→ a*_2_ and *e*_1_ *→ a*_2_. The bridge leads to motif 2, which is the following cycle:

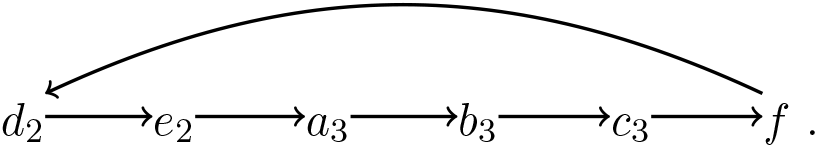

The bridge transitions to motif 2 via *c*_2_ *→ d*_2_. There is an additional state for *b*, which follows from the Start-state; this state transitions to the bridge via *b*_4_ *→ a*_2_, and to motif 2 via *b*_2_ *→ c*_3_. The consequences of pairwise merging of states are shown in Table S2 (Supporting Information). The mergers create unobserved subsequences such as *b → a → ω* (type I context dependence) or significantly enhance probabilities of subsequences such as *b → a → b → c → f* (type II context dependence).

After deafening, a novel transition *d → a* appears. Moreover, the probability of stopping after syllable *a* is strongly enhanced. State multiplicity disappears except for syllable *b*, which is still associated with two states, reflecting the context-dependent transitions

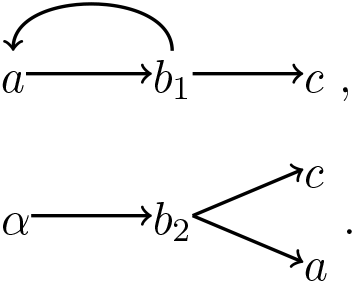

This is a type II context dependence. In both cases, syllable *b* is followed by syllable *a* or *c*. However, from *b*_1_ the transition to *c* is favored, with probability 0.9; in contrast, from *b*_2_ the transition to *a* is favored with probability 0.8. Merging *b*_1_ and *b*_2_ enhances the subsequence *α → b → c* (observed *P*_*s*_ = 0.2) to median *P*_*s*_ = 0.5, which is significant (*p* = 0).

### o10bk90

In the normal condition, syllables *f* and *g* are represented by two states each (Fig. 4), reflecting the following context dependencies in syllable transitions:

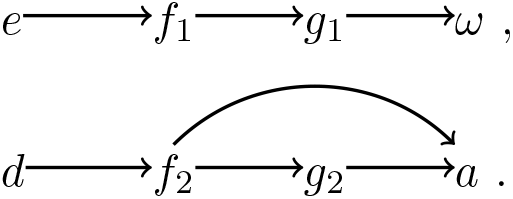

Merging *f*_1_ and *f*_2_ creates an unobserved subsequence *e → f → a* with median *P*_*s*_ = 0.1 and *p* = 0.0002. Merging *g*_1_ and *g*_2_ enhances the probability of the subsequence *d → f → g → ω* from *P*_*s*_ = 0.02 to median *P*_*s*_ = 0.3, which is significant (*p* = 0).

After deafening, transition *α → d* is weakened, and transitions *α → a* and *α → g* become stronger (Fig. 4). The state multiplicity for *f* persists, reflecting the context-dependent transitions

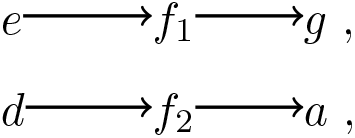

which is the same as in the normal condition. As in the normal condition, merging *f*_1_ and *f*_2_ creates the unobserved subsequence *e → f → a* with median *P*_*s*_ = 0.1 and *p* = 0.

The subsequence *d → f → g* becomes rare after deafening (*P*_*s*_ = 0.5, before deafening; *P*_*s*_ = 0.007, deafened), indicating that deafening makes the transition *f*_2_ *→ g*_2_ rare. Syllable *g* is now represented with one state only, because this does not make the subsequence *d → f → g → ω* more frequent than observed, unlike in the normal condition.

## Supplementary Tables

**Table S1.** Consequences of pairwise merging of states with the same syllables in the POMM for bfa7 in the normal condition. Listed are the pair of states merged, subsequences examined, *P*_*s*_ of the subsequences in the observed set, median of the *P*_*s*_ distribution generated from the state-merged POMMs, and the p-value.

| states merged | subsequence | $P_s$ observed | median $P_s$ after merging | $p$ |
| --- | --- | --- | --- | --- |
| $b_1, b_2$ | *abcdg* | 0 | 0.07 | 0.004 |
| $c_1, c_2$ | *fc dg* | 0 | 0.1 | 0 |
| $c_1, c_3$ | *bcdcdb* | 0 | 0.04 | 0.01 |
| $c_1, c_4$ | *fcdb* | 0 | 0.08 | 0 |
| $c_2, c_3$ | *abcdg* | 0 | 0.04 | 0.007 |
| $c_2, c_4$ | *bcd b* | 0 | 0.04 | 0.003 |
| $c_3, c_4$ | *bcd b* | 0 | 0.04 | 0.009 |
| $d_1, d_2$ | *fc dg* | 0 | 0.1 | 0.0001 |
| $d_1, d_3$ | *bcdcdb* | 0 | 0.04 | 0.01 |
| $d_1, d_4$ | *fcdb* | 0 | 0.08 | 0.0001 |
| $d_2, d_3$ | *abcdg* | 0 | 0.04 | 0.006 |
| $d_2, d_4$ | *bcd b* | 0 | 0.04 | 0.0066 |
| $d_3, d_4$ | *bcd b* | 0 | 0.04 | 0.009 |

**Table S2.**
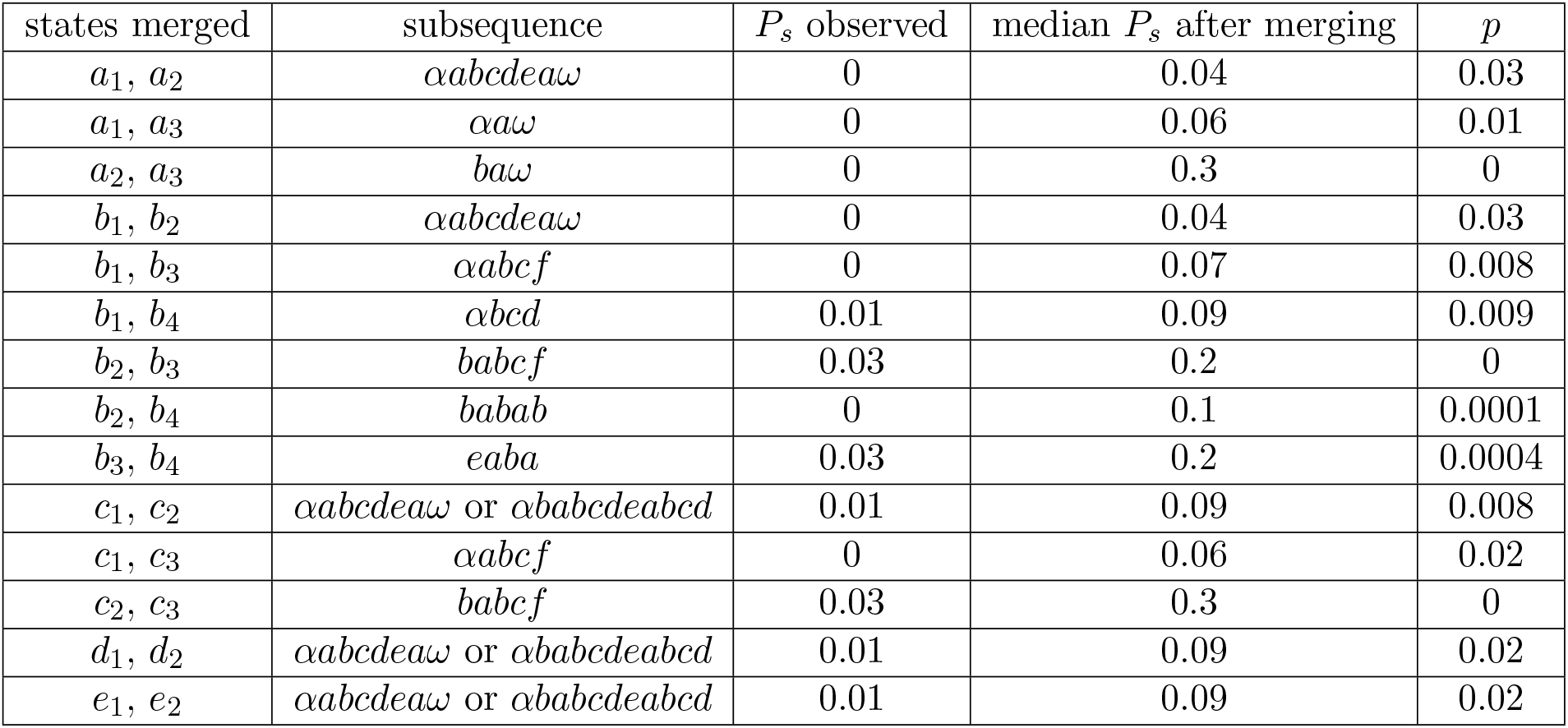
Consequences of pairwise merging of states with the same syllables for o46bk78 in the normal condition.

## Supplementary Figures

**Fig. S1.**
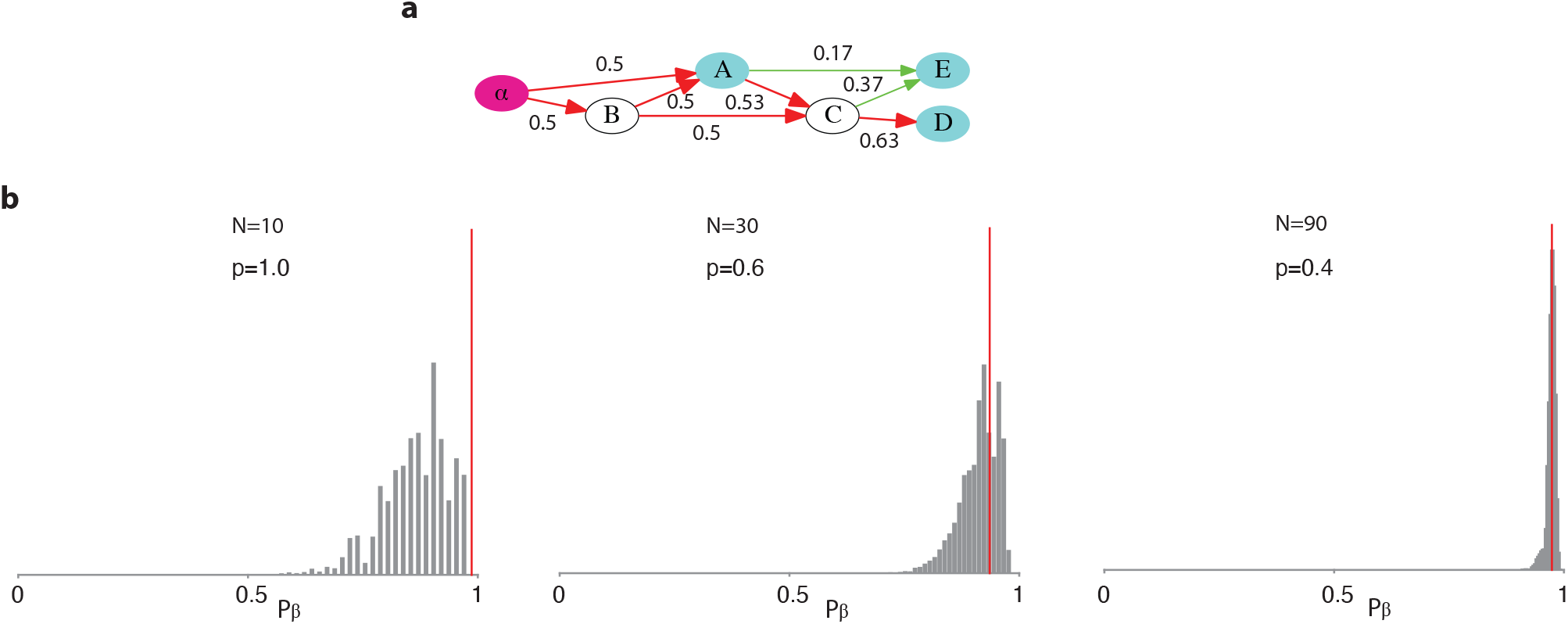
Statistical tests of the Markov models. **a**. The ground-truth model is a Markov model. The transition probabilities are marked near the arrows. **b**. Statistical tests of the Markov models for *N* = 10, 30, 90. The gray bars are the distributions of *P*_*β*_ for the generated sets. The red lines are the *P*_*β*_ for the observed sets. The Markov model is accepted in all cases.

**Fig. S2.**
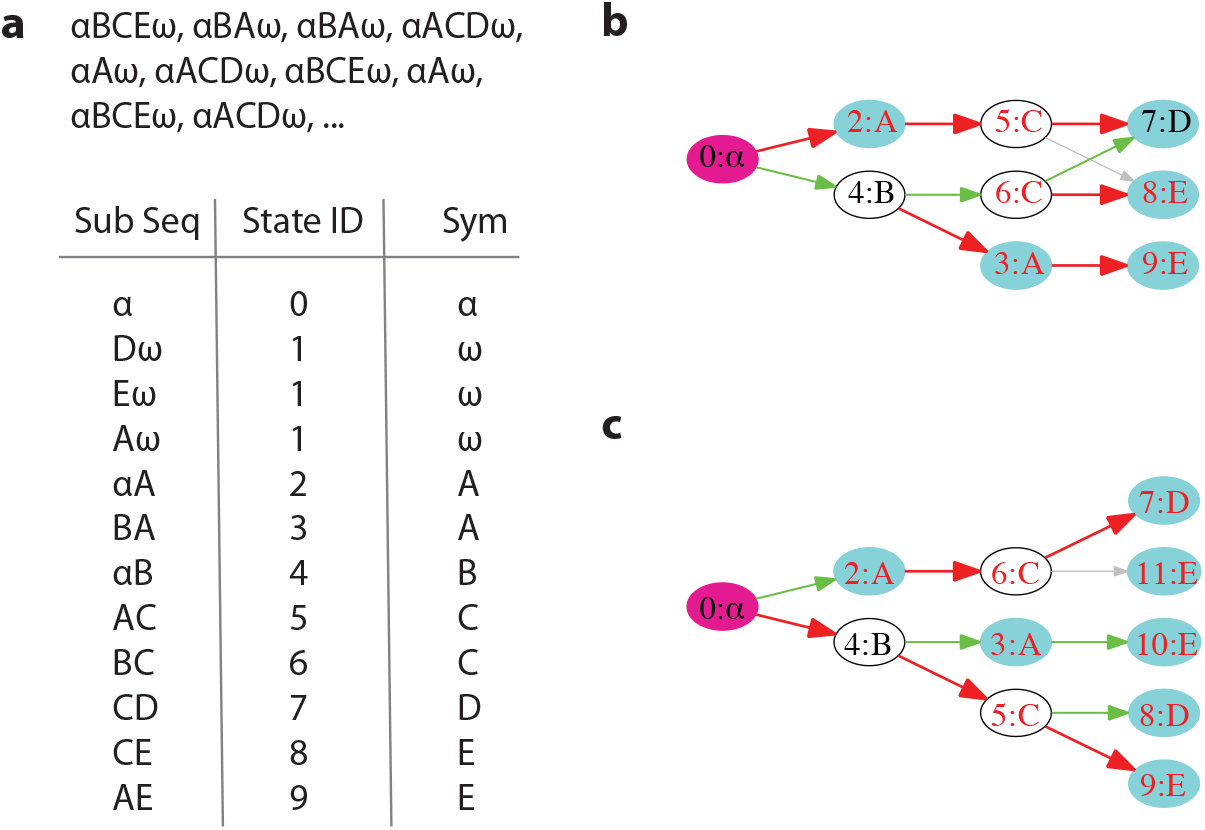
Constructing POMMs that are equivalent to higher order Markov models. An example is shown for constructing POMMs from the second and third order Markov models. The observed sequences (*N* = 30) are generated from the ground truth POMM in Fig. 2a. **a**. The process of assigning states for the second order Markov model. Examples of sequences are shown. Each sequence is flanked by the start symbol *α* and the end symbol *ω*. In the table, the first column shows the unique subsequences. The second column shows the state IDs assigned to the subsequences. The third column shows the syllables (or symbols) associated with the states. **b**. The POMM corresponding to the second order Markov model. Both the state IDs and the associated syllables (or symbols) are displayed. **c**. The POMM corresponding to the third order Markov model. Both models have *p* = 1, hence they are compatible with the observed sequences. The POMMs have more extra states than the ground truth model, which has two states for *A* and two states for *C*.

**Fig. S3.**
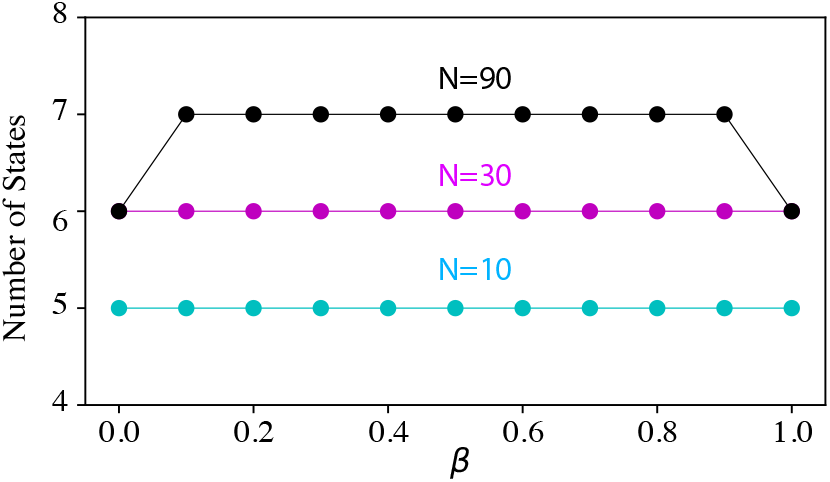
The impact of the choice of *β* in inferring POMMs. The median number of states in the POMMs, inferred from the observed sequences generated from the ground truth model (Fig. 2a), is presented for various values of *β*. The sample sizes considered are *N* = 10, 30, 90. The inferred models exhibit consistent structures across a wide range of *β* values.

**Fig. S4.**
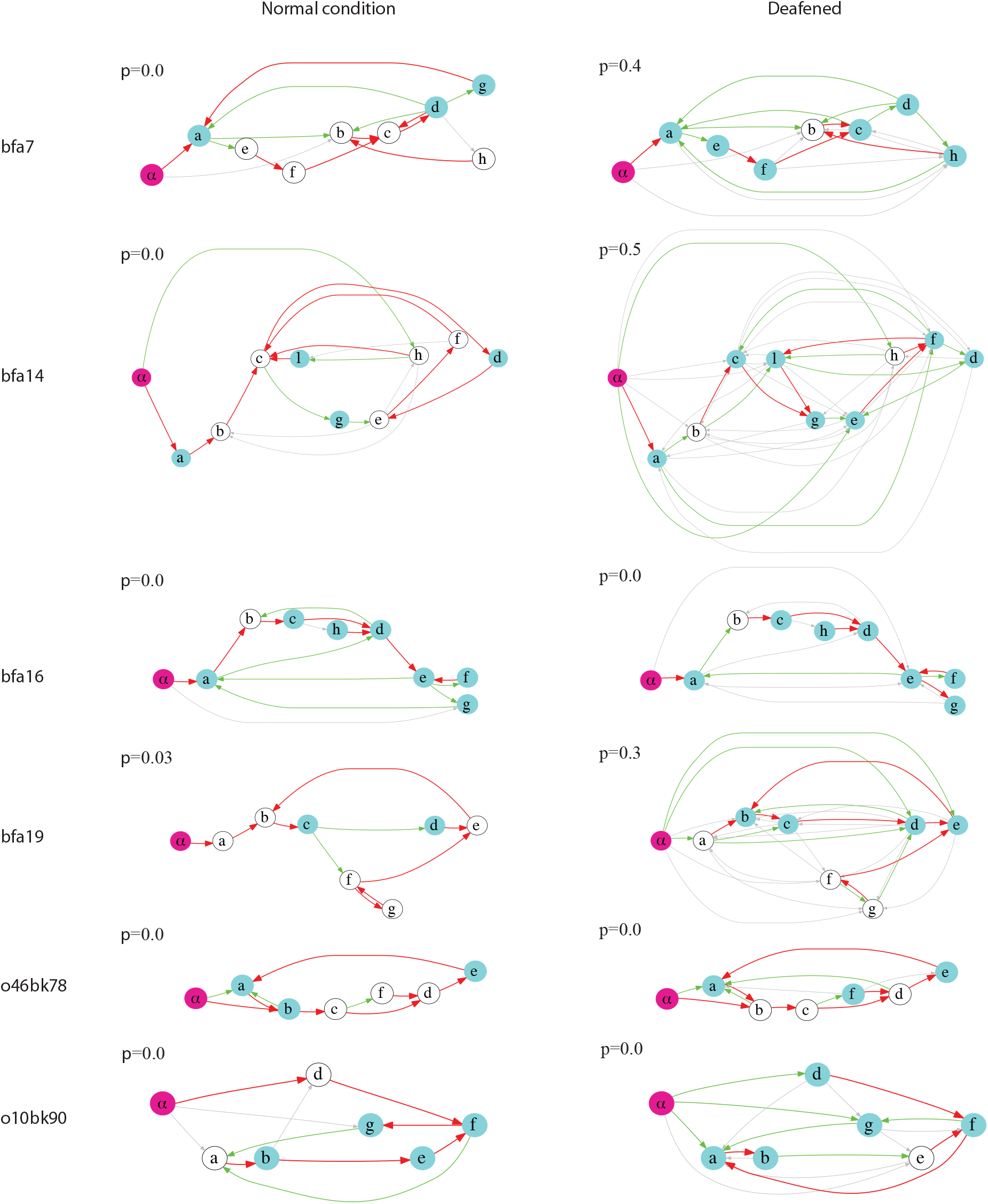
Markov models for all birds. Bird names and the p-values are displayed.

**Fig. S5.**
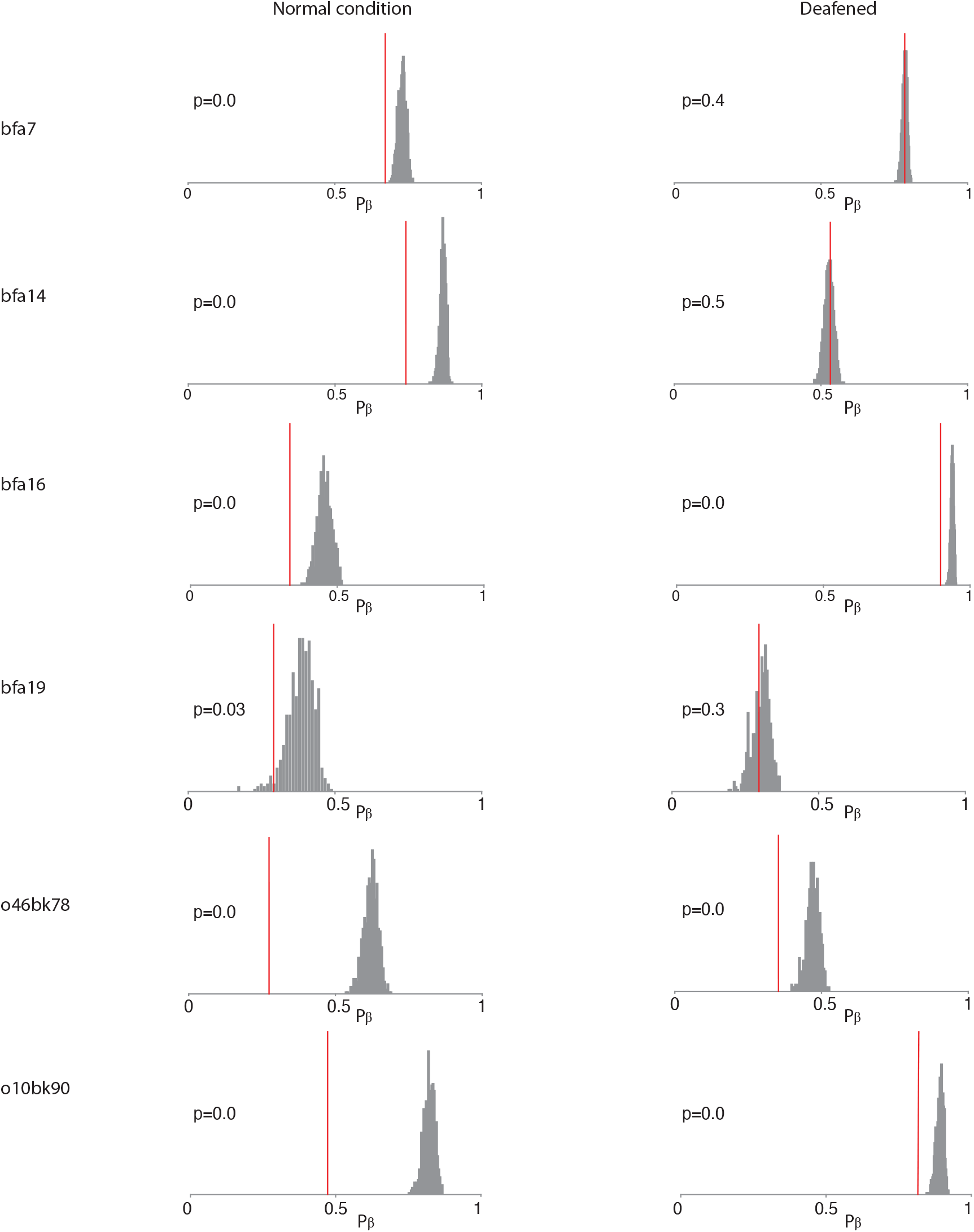
Statistical tests of the Markov models for all birds. The distributions of *P*_*β*_ for the generated sets and *P*_*β*_ of the observed set (red line) are shown. The p-values are displayed.

**Fig. S6.**
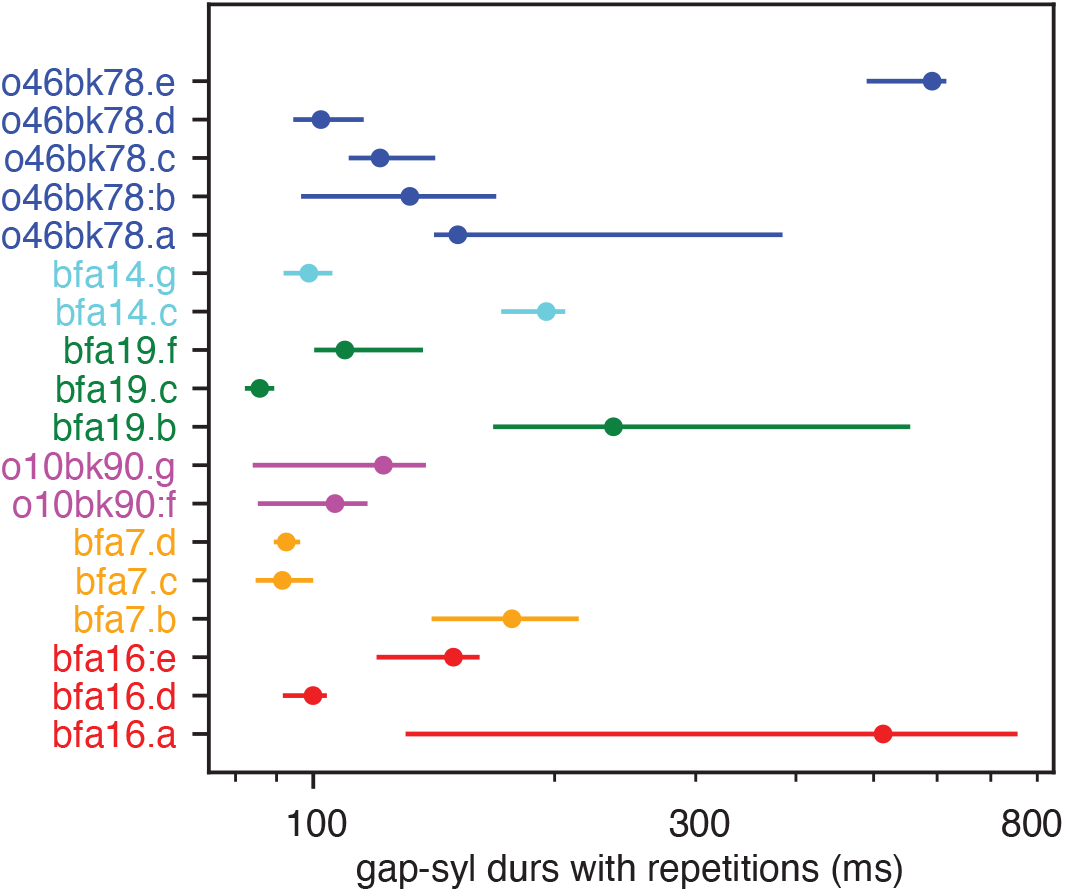
Durations of gap-syllables including repetitions. If a syllable repeats, the duration is defined as the period from the onset of the repetition to the offset of the repetition plus the gap preceding the onset.

**Fig. S7.**
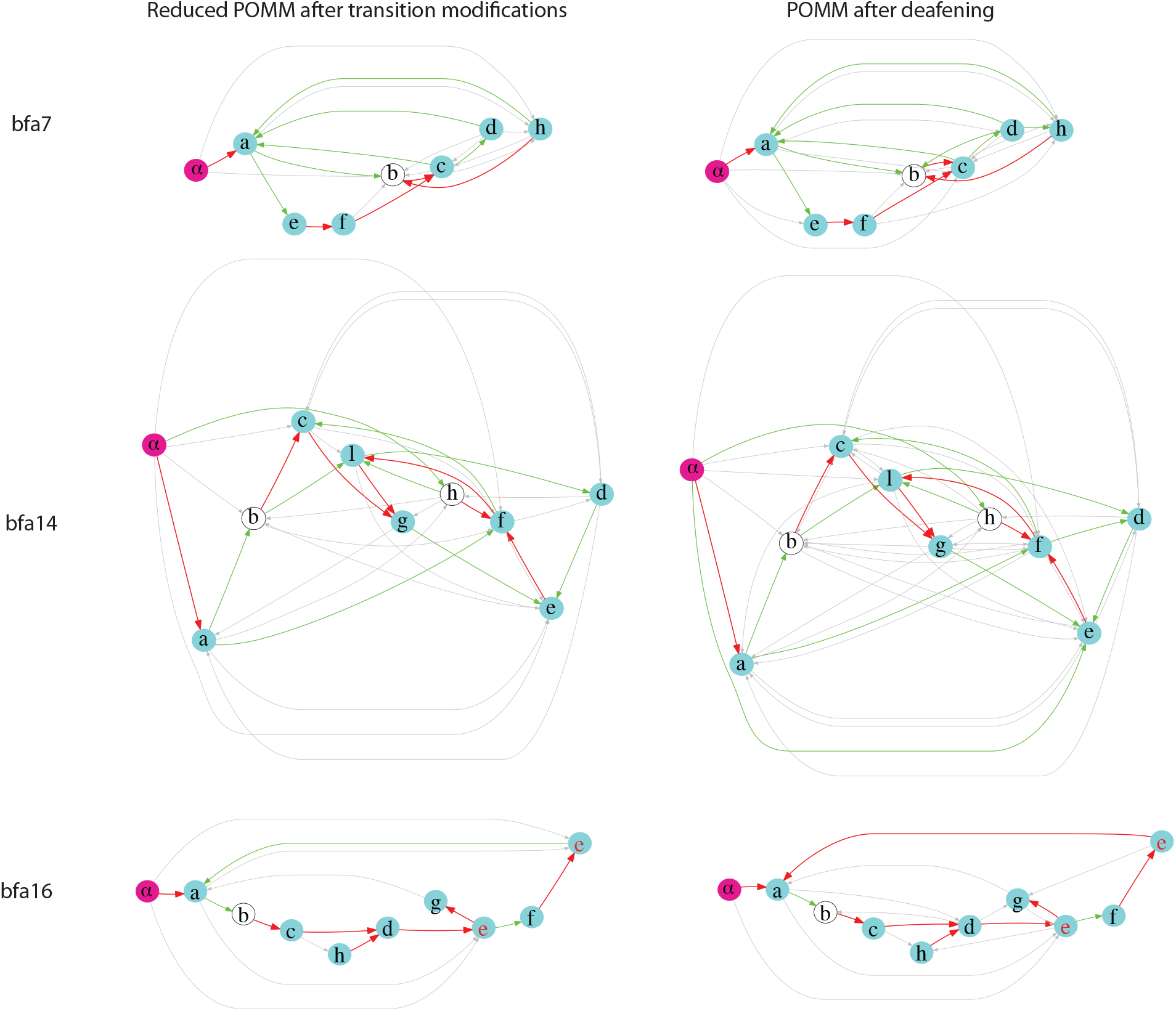
Comparisons of the reduced POMMs and the POMMs after deafening for three birds. The results for bfa7, bfa14, and bfa16 are shown.

**Fig. S8.**
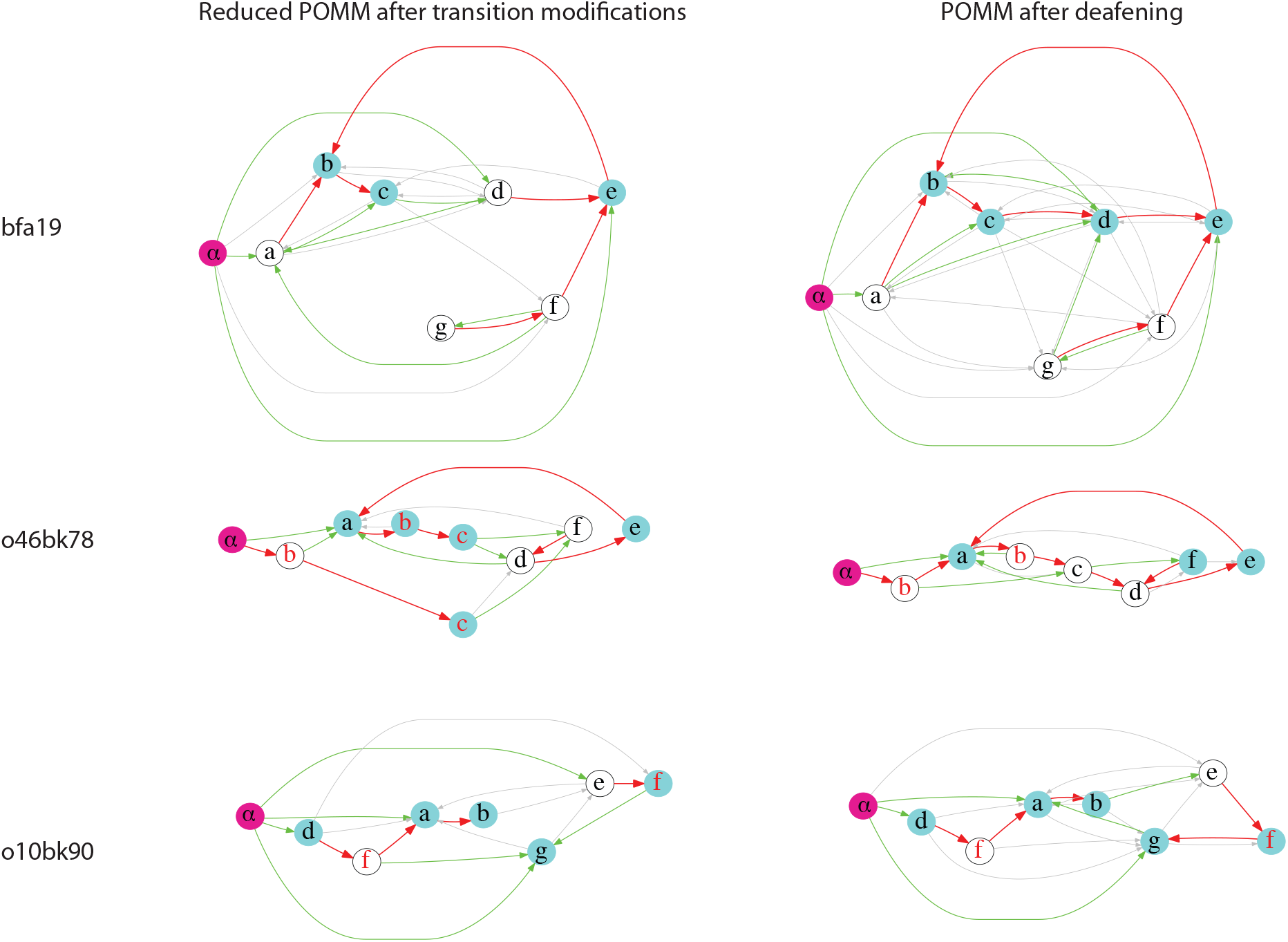
Comparisons of the reduced POMMs and the POMMs after deafening for three birds. The results for bfa19, o46bk78, and o10bk90 are shown.

**Fig. S9.**
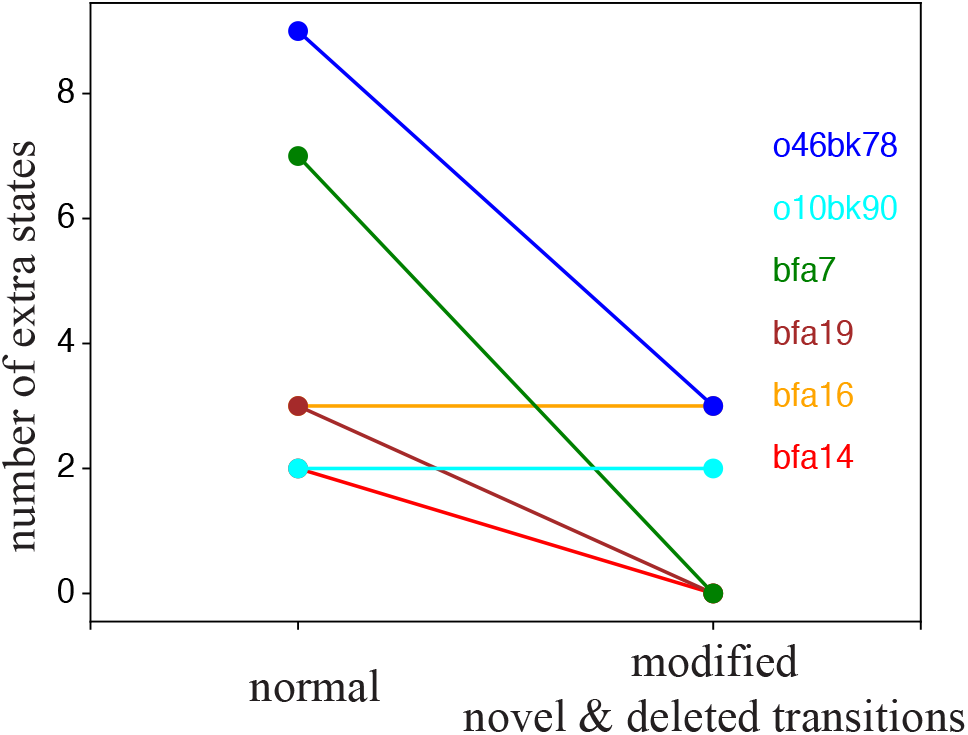
Changes in the number of extra states in the reduced POMMs. The transition probabilities are modified by solely adding novel transitions and deleting dropped ones. For bfa7, bfa19, and bfa14, the reduced POMMs become Markovian. In the case of o46bk78, there is a reduction in state multiplicity compared to the POMM under the normal condition. No change in state multiplicity is observed for bfa16 and o10bk90.

**Fig. S10.**
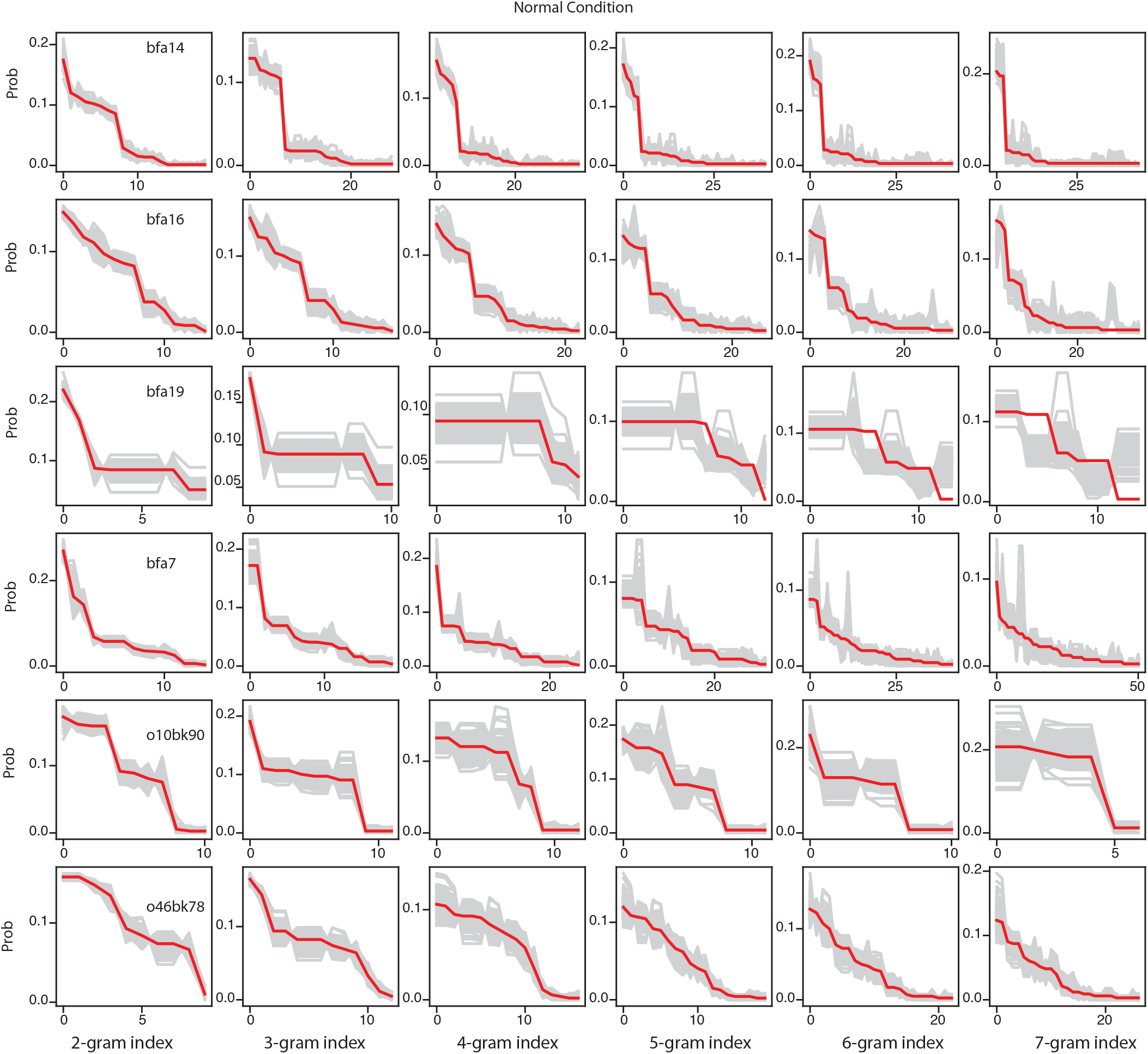
Comparisons of n-gram distributions in the normal condition. The redlines are the probabilities of n-grams in the observed sets. The n-grams are ordered in a descending order of probabilities in the observed sets. The gray lines are the probabilities of the ordered n-grams computed from 100 sets of sequences generated from the POMMs. The red line is mostly within the range defined by the gray lines.

**Fig. S11.**
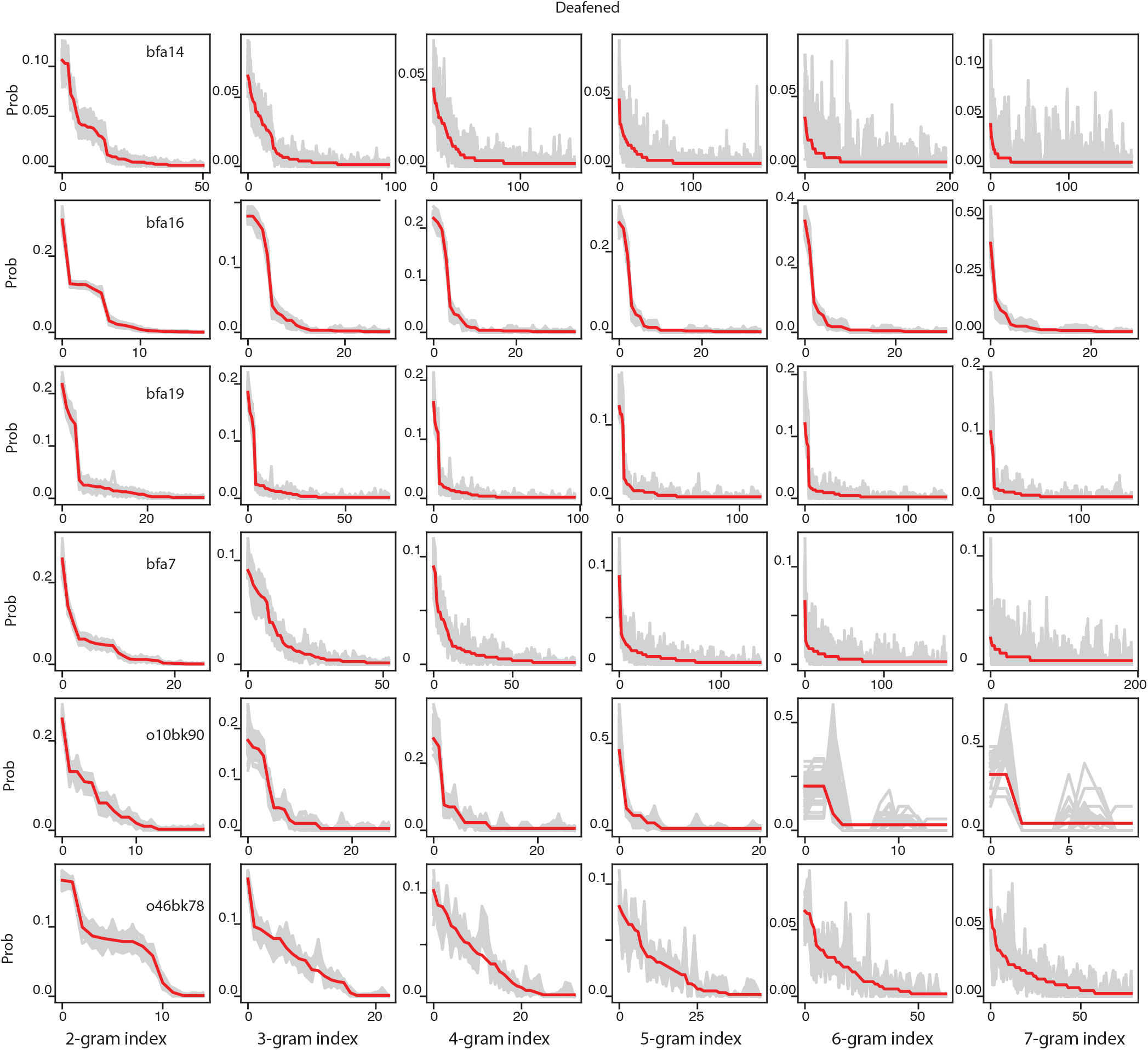
Comparisons of n-gram distributions in after deafening. The same as in Fig. S9 for the deafened case.

**Fig. S12.**
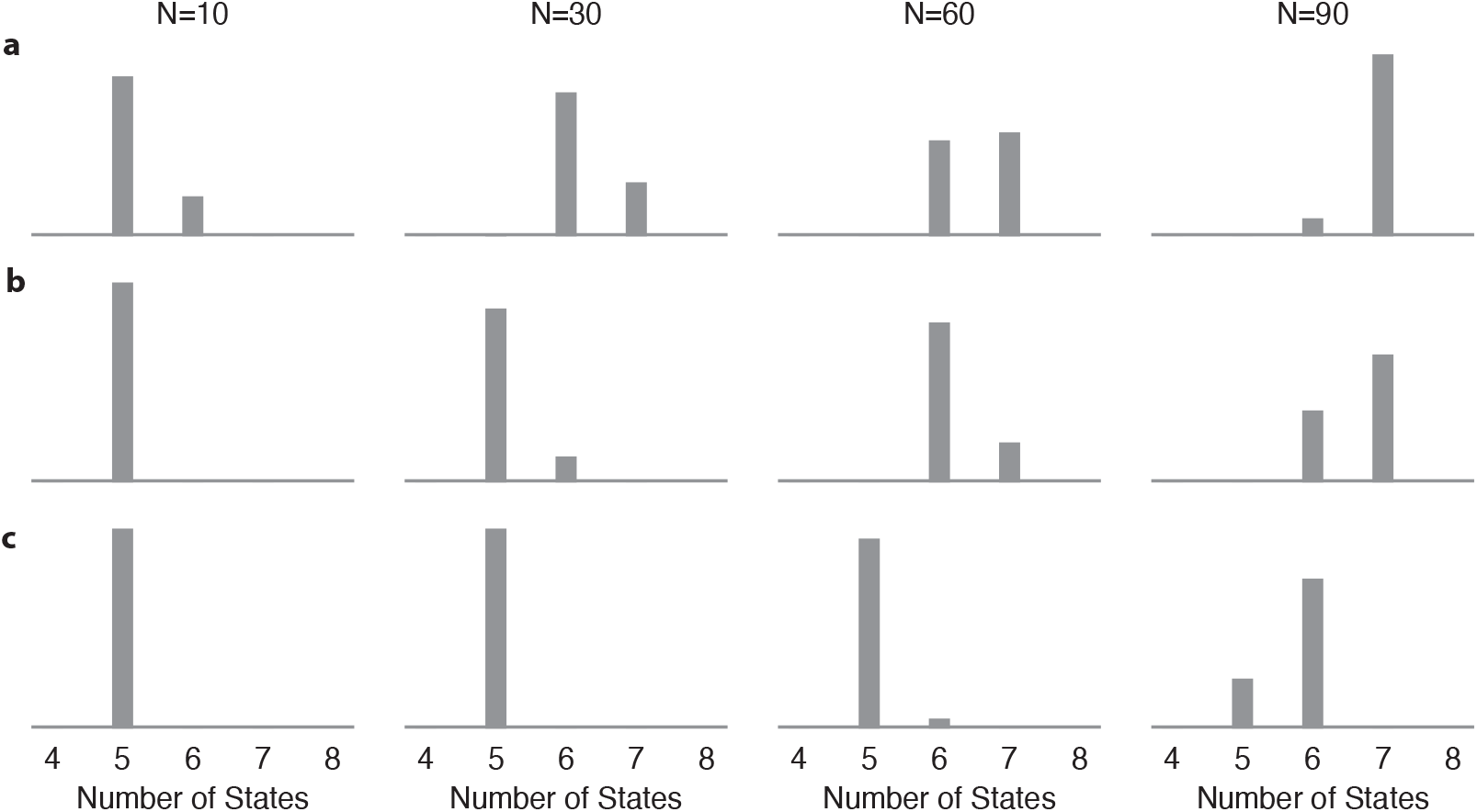
Comparisons POMMs inferred using our method, AIC and BIC. *N* “observed sequences” are generated with the ground truth model shown in Fig. 2a, which has 7 states excluding the start state and the end state. The histograms of the number of states for the syllables in the inferred POMMs in 100 runs are shown for (a) our method; (b) AIC; (c) BIC. The minimal models for AIC and BIC are selected among all POMMs with at most two states for each syllable. In all cases of *N* = 10, 30, 60, 90, the AIC and BIC under estimates the number of states compared to our method.

## Notes

Conflict of Interest: The authors declare no competing financial interests.

### Competing Interest Statement

The authors have declared no competing interest.

### Summary of Updates

The manuscript has been revised to enhance clarity in its presentation. We have made efforts to improve readability, without altering the content from the previous versions. This revision aims to facilitate a better understanding of the material.

## References

Bialek W, Nemenman I, Tishby N (2001) Predictability, complexity, and learning. Neural computation 13:2409–2463.

Cohen Y, Shen J, Semu D, Leman DP, Liberti WA, Perkins LN, Liberti DC, Kotton DN, Gardner TJ (2020) Hidden neural states underlie canary song syntax. Nature 582:539–544.

Egger R, Tupikov Y, Elmaleh M, Katlowitz KA, Benezra SE, Picardo MA, Moll F, Kornfeld J, Jin DZ, Long MA (2020) Local axonal conduction shapes the spatiotemporal properties of neural sequences. Cell 183:537–548.

Ellson J, Gansner E, Koutsofios L, North SC, Woodhull G (2001) Graphviz – open source graph drawing tools In International Symposium on Graph Drawing, pp. 483–484. Springer.

Fee MS, Kozhevnikov AA, Hahnloser RH (2004) Neural mechanisms of vocal sequence gener-ation in the songbird. Annals of the New York Academy of Sciences 1016:153–170.

Gibbs AL, Su FE (2002) On choosing and bounding probability metrics. International statistical review 70:419–435.

Hahnloser RH, Kozhevnikov AA, Fee MS (2002) An ultra-sparse code underlies the generation of neural sequences in a songbird. Nature 419:65.

Hanuschkin A, Diesmann M, Morrison A (2011) A reafferent and feed-forward model of song syntax generation in the bengalese finch. Journal of computational neuroscience 31:509–532.

Isola GR, Vochin A, Sakata JT (2020) Manipulations of inhibition in cortical circuitry differentially affect spectral and temporal features of bengalese finch song. Journal of Neurophysiology 123:815–830.

Jin DZ (2009) Generating variable birdsong syllable sequences with branching chain networks in avian premotor nucleus hvc. Physical Review E 80:051902.

Jin DZ (2013) The neural basis of birdsong syntax. Progress in cognitive science: From cellular mechanisms to computational theories .

Jin DZ, Kozhevnikov AA (2011) A compact statistical model of the song syntax in bengalese finch. PLoS computational biology 7:e1001108.

Jin DZ, Ramazanoğlu FM, Seung HS (2007) Intrinsic bursting enhances the robustness of a neural network model of sequence generation by avian brain area hvc. Journal of computational neuroscience 23:283–299.

Jun JK, Jin DZ (2007) Development of neural circuitry for precise temporal sequences through spontaneous activity, axon remodeling, and synaptic plasticity. PLoS One 2:e723.

Long MA, Jin DZ, Fee MS (2010) Support for a synaptic chain model of neuronal sequence generation. Nature 468:394.

Lynch GF, Okubo TS, Hanuschkin A, Hahnloser RH, Fee MS (2016) Rhythmic continuous-time coding in the songbird analog of vocal motor cortex. Neuron 90:877–892.

Markowitz JE, Ivie E, Kligler L, Gardner TJ (2013) Long-range order in canary song. PLoS computational biology 9:e1003052.

Nottebohm F, Stokes TM, Leonard CM (1976) Central control of song in the canary, serinus canarius. Journal of Comparative Neurology 165:457–486.

Okanoya K (2004) The bengalese finch: a window on the behavioral neurobiology of birdsong syntax. Annals of the New York Academy of Sciences 1016:724–735.

Okanoya K, Yamaguchi A (1997) Adult bengalese finches (lonchura striata var. domestica) require real-time auditory feedback to produce normal song syntax. Journal of neurobiology 33:343–356.

OpenAI (2023) Gpt-4 technical report.

Picardo MA, Merel J, Katlowitz KA, Vallentin D, Okobi DE, Benezra SE, Clary RC, Pnevmatikakis EA, Paninski L, Long MA (2016) Population-level representation of a temporal sequence underlying song production in the zebra finch. Neuron 90:866–876.

Rabiner LR (1989) A tutorial on hidden markov models and selected applications in speech recognition. Proceedings of the IEEE 77:257–286.

Sakata JT, Brainard MS (2006) Real-time contributions of auditory feedback to avian vocal motor control. Journal of Neuroscience 26:9619–9628.

Sakata JT, Brainard MS (2008) Online contributions of auditory feedback to neural activity in avian song control circuitry. Journal of Neuroscience 28:11378–11390.

Schmidt MF (2003) Pattern of interhemispheric synchronization in hvc during singing correlates with key transitions in the song pattern. Journal of neurophysiology 90:3931–3949.

Tupikov Y, Jin DZ (2021) Addition of new neurons and the emergence of a local neural circuit for precise timing. PLoS computational biology 17:e1008824.

Virtanen P, Gommers R, Oliphant TE, Haberland M, Reddy T, Cournapeau D, Burovski E, Peterson P, Weckesser W, Bright J, van der Walt SJ, Brett M, Wilson J, Millman KJ, Mayorov N, Nelson ARJ, Jones E, Kern R, Larson E, Carey CJ, Polat İ, Feng Y, Moore EW, VanderPlas J, Laxalde D, Perktold J, Cimrman R, Henriksen I, Quintero EA, Harris CR, Archibald AM, Ribeiro AH, Pedregosa F, van Mulbregt P, SciPy 1.0 Contributors (2020) SciPy 1.0: Fundamental Algorithms for Scientific Computing in Python. Nature Methods 17:261–272.

Wittenbach JD, Bouchard KE, Brainard MS, Jin DZ (2015) An adapting auditory-motor feedback loop can contribute to generating vocal repetition. PLoS computational biology 11:e1004471.

Woolley SM, Rubel EW (1997) Bengalese finches lonchura striata domestica depend upon auditory feedback for the maintenance of adult song. Journal of Neuroscience 17:6380–6390.

Woolley SM, Rubel EW (2002) Vocal memory and learning in adult bengalese finches with regenerated hair cells. Journal of Neuroscience 22:7774–7787.

Zhang YS, Wittenbach JD, Jin DZ, Kozhevnikov AA (2017) Temperature manipulation in songbird brain implicates the premotor nucleus hvc in birdsong syntax. Journal of Neuroscience 37:2600–2611.

Zucchini W, MacDonald IL (2009) Hidden Markov models for time series: an introduction using R Chapman and Hall/CRC.

